# A geometric parameterization for beta turns

**DOI:** 10.1101/2024.01.01.573818

**Authors:** Nicholas E. Newell

## Abstract

Beta turns, in which the protein backbone abruptly changes direction over four amino acid residues, are the most common type of protein secondary structure after alpha helices and beta sheets and play many key structural and functional roles. Previous work has produced classification systems for turn backbone geometry at multiple levels of precision, but these all operate in backbone dihedral-angle (Ramachandran) space, and the absence of a local Euclidean-space coordinate system and structural alignment for turns, or of any systematic Euclidean-space characterization of turn backbone shape, presents challenges for the visualization, comparison and analysis of the wide range of turn conformations and the design of turns and the structures that incorporate them. This work derives a local coordinate system that implicitly aligns turns, together with a simple geometric parameterization for turn backbone shape that describes modes of structural variation not explicitly captured by existing systems. These modes are shown to be meaningful by the demonstration of clear relationships between parameter values and the electrostatic energy of the beta-turn H-bond, the overrepresentation of key side-chain motifs, and the structural contexts of turns. Geometric turn parameters, which complement existing Ramachandran-space classifications, can be used to tune turn structures for compatibility with particular side-chain interactions or contexts, and they should prove valuable in applications, such as protein design, where an enhanced Euclidean-space description of turns may improve understanding or performance. The web-based tools ExploreTurns, MapTurns and ProfileTurn, available at www.betaturn.com, incorporate turn-local coordinates and turn parameters and demonstrate their utility.

## 1 Introduction

### 1.1 Background

Abrupt changes in the direction of the protein backbone (BB) chain are accomplished by tight turns of types {δ, γ, β, α, π}, with lengths of {2, 3, 4, 5, 6} amino acid (AA) residues^1^. Of these types, the four-residue beta turn is found most frequently; it constitutes the most common protein secondary structure after alpha helices and beta strands. Either individually or in overlapping multiples^2^, beta turns are found in loops that link segments of repetitive BB structure, and they are also very common in extended loops. Turns exhibit a wide range of BB conformations which, taken together with the variety of intra- and extra-turn structural interactions that their side-chains (SCs) can mediate, enable them to play crucial roles in multiple contexts in proteins; for an overview, see *Roles of beta turns in proteins* at www.betaturn.com.

Beta turns were first described in 1968 by Venkatachalam^3^, who determined the allowed geometries of four-residue chain segments that exhibit a 4>1 H-bond. Since this study, a substantial body of analytical work (see short reviews in^4,5^) has evolved the beta-turn definition and identified and classified new geometries. A current definition^4^, applied here, specifies a four-residue BB segment in which the distance between the alpha carbons of the first and last residues is no greater than 7Å and the two central residues are not part of helices or strands. In about half of the segments that meet this definition, the BB NH group of the fourth turn residue (MCN4) donates an H-bond to the BB carbonyl oxygen of the first residue (MCO1), and the presence of this 4>1 H-bond by itself constitutes an alternative, more restrictive definition.

Three beta-turn classification systems developed in previous studies are employed in this work; each partitions turns by BB geometry in Ramachandran space at a different level of precision. These systems, summarized in the ExploreTurns^6^ online help at www.betaturn.com, are referred to here, in order of increasing precision, as the “classical” type system^7,8,9,10^ and the BB cluster and Ramachandran type systems^4^, with the last system based on the map of Ramachandran space due to Hollingsworth and Karplus^11^.

### 1.2 Motivation and description of the present work

The inherent difficulty of visualizing and contrasting the wide range of beta-turn geometries in Ramachandran space^12^ motivated the construction of a turn-local coordinate system which could provide a common Euclidean-space framework for structural analysis, and the derivation of geometric parameters for turns was suggested by the observation that while the shape of a protein helix, which can be described as a series of overlapping turns, has been characterized by parameters such as twist, rise, and pitch that vary between helix types and also with the irregularities of helices within each type^13^, no such parameterization had been developed to describe the bulk BB shapes of beta turns, which vary between and within turn types or BB clusters.

The case for the development of geometric parameters was bolstered by the need for structural discriminators within types and clusters: the set of turns with both central residues in or near the right-handed helical conformation, which constitutes close to half of all turns, is represented by a single classical type (I) and a single BB cluster (with Ramachandran-type label AD)^4^, yet, as shown below, type I/cluster AD includes a range of conformations which are compatible to widely varying degrees with key SC motifs, or compatible with different SC motifs entirely, and can play distinct roles in the structural contexts that incorporate beta turns. The presence of a meaningful range of BB structures within type I/cluster AD, and also, to varying degrees, within other, smaller types/clusters, prompted a search for a set of parameters capable of characterizing modes of geometric variation not explicitly captured by Ramachandran-space systems.

The construction of a turn-local coordinate system is achieved via a set of geometric definitions, and the representation of turns in the local system implicitly aligns them. Using the definitions, the turn parameters are then derived, their modes of variation are demonstrated with example structures, and their distributions within each classical type and BB cluster are characterized. Parameter distributions are also heat-mapped across the Ramachandran-angle spaces of the two central turn residues, revealing the bond rotations that drive parameter variation in each type or cluster.

The fractional overrepresentation of key sequence motifs in turns is shown to vary dramatically with parameter values, demonstrating the utility of parameters as structural discriminators that can support geometric tuning for compatibility with SC interactions. Relationships are also demonstrated between parameter values and the electrostatic energy of the 4>1 H-bond.

The roles and contexts of parameter regimes are explored, as are the effects of *cis*-/cis-Pro-peptide bonds on parameter values. The dependencies between parameters are evaluated, and the potential utility of parameters in structure validation is investigated.

Many of the results presented here were generated with ExploreTurns^6^, a web facility for the comprehensive exploration, analysis, geometric tuning and retrieval of beta turns and their contexts from the dataset compiled for this study. A second tool, MapTurns^14^, produces interactive, 3D conformational heatmaps of the BB and SC structure, H-bonding and contexts associated with sequence motifs in beta turns, and a third facility, ProfileTurn, profiles a turn uploaded by the user and evaluates the compatibility between its geometry and sequence motif content. Extensive online support is available within all three tools, which are available at www.betaturn.com.

## 2 Methods

### 2.1 Euclidean-space representation and alignment

The derivation of turn parameters begins with a set of geometric definitions which establish a common turn-local Euclidean-space coordinate system. The **span line** is the line between a turn’s first and last alpha carbons (*C*_α1_and *C*_α4_), and the span, already used in the beta-turn definition, is its length. The **turn center** is the midpoint of (*C*_2_→*N*_3_), the middle peptide bond in the turn. The **turn plane** contains the span line and passes through the turn center, and the perpendicular dropped from the turn center to the span line defines the **turn axis**. Using these definitions, a local orthogonal right-handed coordinate system is established. The system’s origin lies at the turn center, while the x axis lies in the turn plane, collinear with the turn axis, with its positive sense oriented from the center towards the span line. The y axis lies in the turn plane, parallel to the span, with its positive sense oriented towards the C-terminal half of the turn. Finally, the z axis is perpendicular to the turn plane, with its positive sense determined by the cross-product of the x and y axes. For a depiction of the turn coordinate system, see any figure below that displays a structure.

The atomic coordinates of each turn are transformed from the global protein system in each turn’s PDB^15^ file to the turn-local system, implicitly aligning the structures. The accuracy of this alignment is tested by a comparison between implicit pairwise turn alignments and the best alignments generated by a perturbative search procedure (see Supplementary Section 1.1).

### 2.2 Parameter definitions

#### Span

The pre-existing span parameter measures the distance [Å] between *C*_α1_ and *C*_α4_, and a 7Å threshold is part of the beta-turn definition. Two new span-related parameters are defined: the N- and C-terminal half-spans measure the distances between the turn axis and *C*_α1_ and *C*_α4_ respectively, and quantify the separate “widths” of the N- and C-terminal halves of the turn, providing an additional degree of freedom in the measurement of span. Span values in the dataset range from a minimum of 3.13Å to the 7Å limit set by the beta-turn definition, while N-terminal half-spans range from 0 to 4.76Å, and C-terminal half-spans from .76Å to 4.82Å.

#### Bulge

Bulge is a measure of the magnitude of the excursion that the BB makes in the turn plane as it bends to form the turn. Bulge is defined as the distance [Å] from the turn center to the span line. Bulge values in the dataset range from 1.28Å to 4.7Å.

#### Skew

Skew is a dimensionless measure of the asymmetry of the projection of a beta turn’s BB onto the turn plane. Skew is defined as the directed displacement along the span line between the line’s center and the point of intersection between the turn axis and the line, divided by half the span. Skew is negative when the axis intersects the span line in its N-terminal half, indicating a “lean” towards the N-terminus, or positive when the axis intersects the line in its C-terminal half, producing a C-terminal lean. Skew is expressed in terms of the half-spans as:

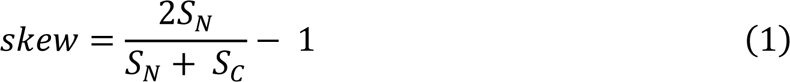

where *S*_*N*_ and *S*_*C*_ are the N- and C-terminal half-spans. Skew is positive when *S*_*N*_ is larger than *S*_*C*_, negative when the reverse is true, or zero when the half-spans are equal. A turn that skews so strongly N-terminally that its axis intersects the span line at the position of *C*_α1_ (*S*_*N*_ = 0) would exhibit a skew of −1, while a turn skewed C-terminally so that its axis intersects the line at *C*_α4_ (*S*_*C*_ = 0) would show a skew of +1. Skew values in the dataset range from −1 to .78.

#### Warp

Warp measures the departure from flatness of a beta turn’s BB in the z dimension of the turn coordinate system, perpendicular to the turn plane. Warp [Å] is defined as the average absolute excursion of the turn’s BB above or below the turn plane between *C*_α1_ and *C*_α4_, excluding BB carbonyl oxygen atoms as well as *C*_α1_ and *C*_α4_ themselves (since they lie within the plane). The eight BB atoms used to compute warp are therefore (C_1_, N_2_, Cα_2_, C_2_, N_3_, Cα_3_, C_3_, N_4_). Warp values in the dataset range from .14Å to 1.25Å. The differences in BB conformation between turn types or BB clusters affect the interpretation and utility of the warp parameter (see Supplementary Section 3.1).

### 2.3 Other methods

Supplementary Sections 1.2-1.4 describe the redundancy-screened beta-turn dataset of 102,192 turns, the H-bond definition, and the statistical models used to calculate the overrepresentation of sequence motifs^16,17^.

## 3 Results

### 3.1 Parameter ranges in structures

Figure 1 shows examples of turns with parameter values near the limits of span, bulge, skew and warp in the dataset.

**Figure 1.**
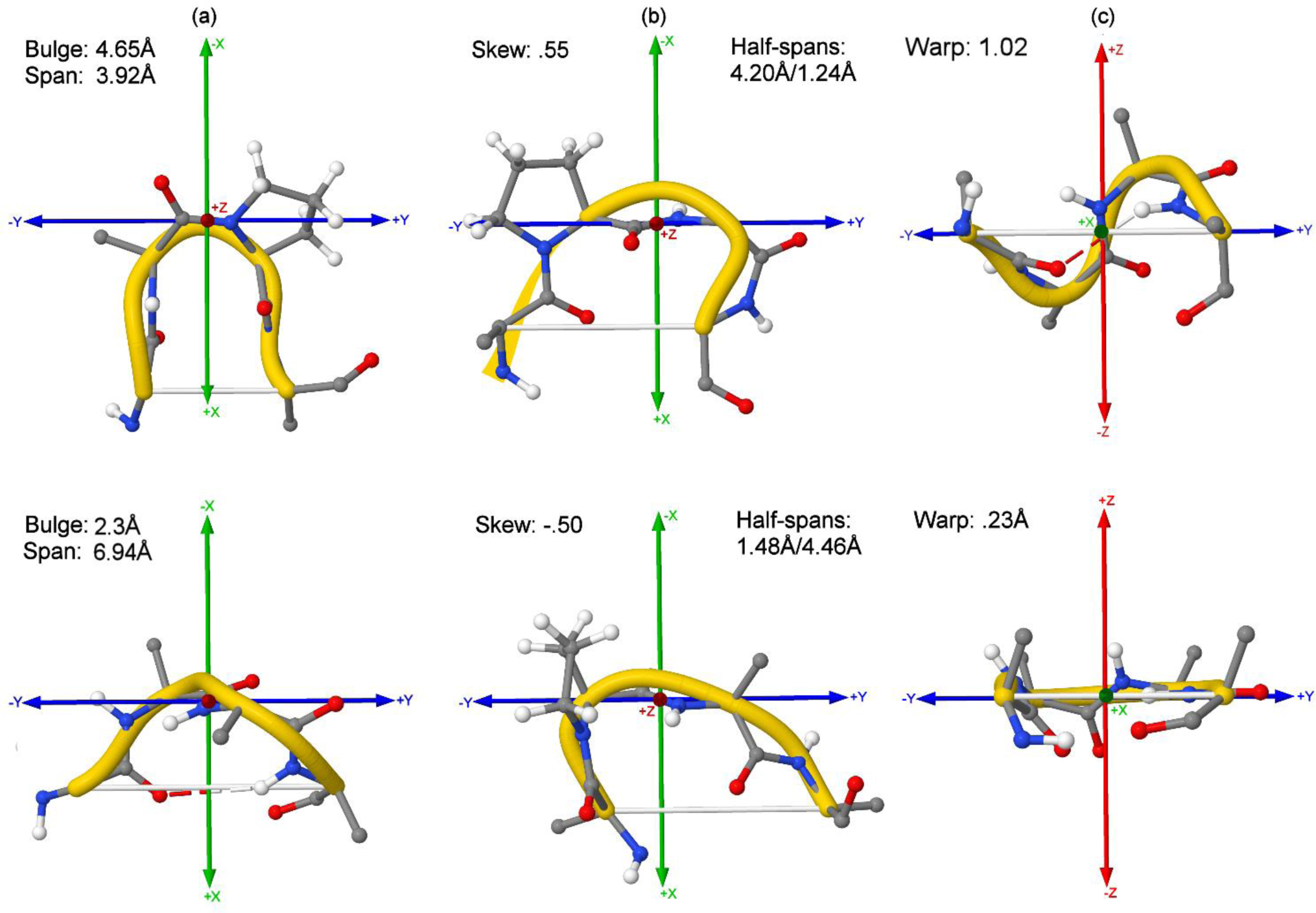
Parameter ranges in structures. Examples of turns that show parameter values near their limits in the dataset. To view any structure in the ExploreTurns tool (available at www.betaturn.com), load the entire database by clicking ***Load Turns Matching Criteria*** without any selection criteria, copy the structure’s address into the ***PDB address*** box, and click ***Browse Structures***. The radius for the display of structure external to the turn is controlled with the ***Display radius*** box; click ***Browse Structures*** again after entering a new radius. While extreme parameter values are correlated with an increased frequency of geometric criteria with outliers in PDB validation checks (Supplementary Section 2.3), none of these examples show outliers. **(a)** 5HKQ_I_74 (top) and 4YZZ_A_239 (bottom) represent the ranges of both bulge and span, since the two parameters exhibit strong negative correlation (Supplementary Section 2.2). **(b)** 5TEE_A_23 (top), and 3NSW_A_44 (bottom) represent strong positive (C-terminal) and negative (N-terminal) skew. Half-spans are also given. **(c)** 5BV8_A_1285 (top) and 3S9D_B_117 (bottom) represent the limits of warp in classical type I, and demonstrate the wide warp range that occurs within this type (and its associated BB cluster, AD_1).

### 3.2 Parameter distributions

Figure 2 plots the distributions of span, bulge, skew and warp in the classical types; these distributions in the BB clusters are plotted in Supplementary Section 2.1. See the figure captions for brief comparative discussions of the distributions, as well as discussions of the partitioning of the distributions in the classical types into those of the BB clusters. Mean and mode parameter values in the BB clusters can be more extreme than those in any classical type; for example, the warp distribution for cluster AG_9 falls to the right of that of any type (Figure S1d), while cluster *cis*DP_18 shows average (negative) skew more than twice as great as any type (Figure S2c).

**Figure 2.**
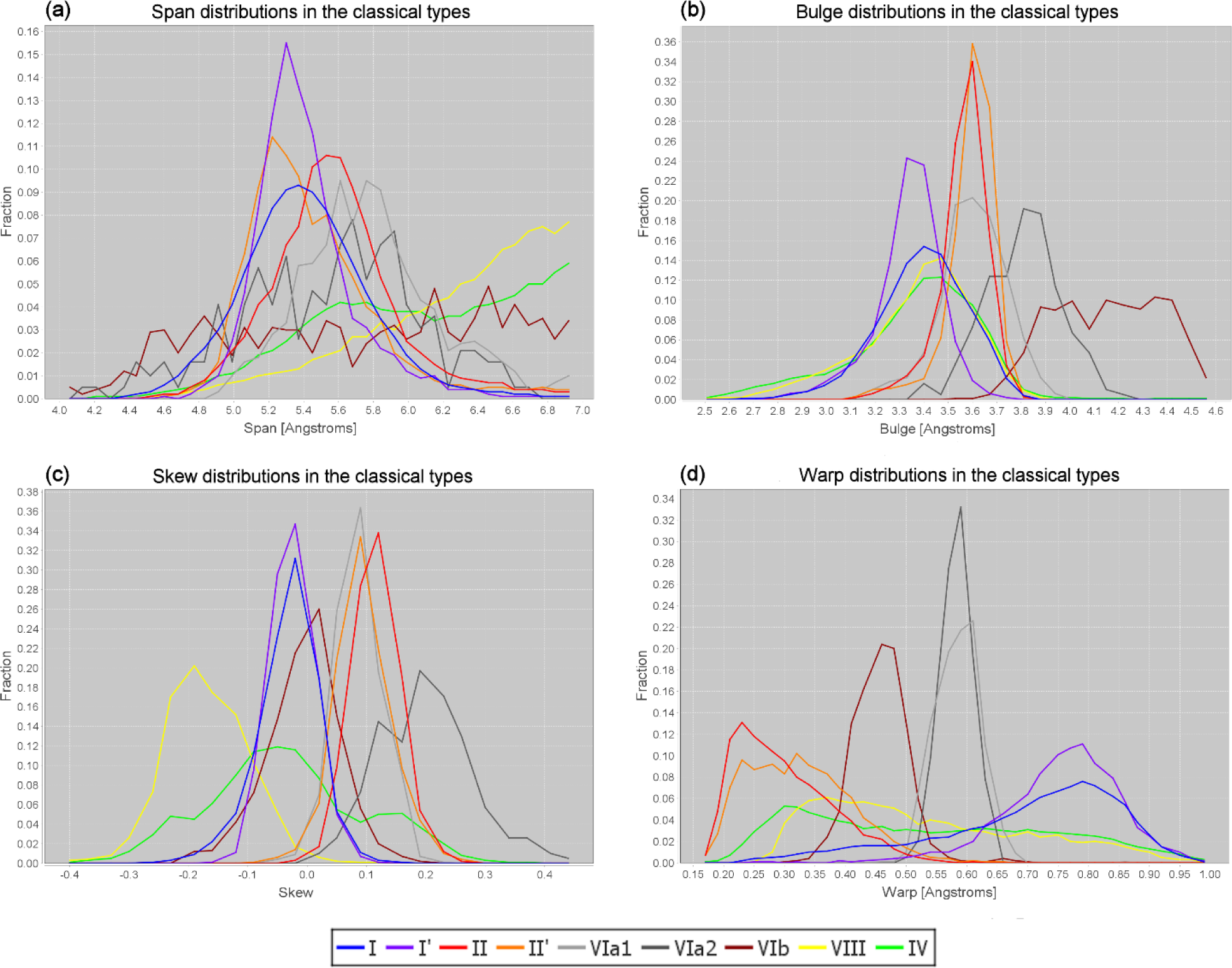
Distributions of the geometric parameters in the classical turn types. Interpolated histograms of the distributions of the geometric parameters in the structurally classified classical turn types and the structurally unclassified type IV. The vertical axis measures turn fraction within each type. **(a)** Span: the type I’ distribution peaks sharply near 5.3Å, reflecting the common role of turns of this type as chain-reversers in 2:2 beta hairpins, and contrasting with the broader peak of the type I distribution, which is consistent with the wider range of type I roles. Types VIb, VIII and IV all show substantial abundance out to the 7Å limit of span, reflecting the scarcity of the constraining 4>1 H-bond in these types. **(b)** Bulge: types VIa2 and VIb extend well to the right of all other types, and types II/II’ show greater bulge than I/I’. All bulge distributions exhibit negative Pearson skewness^18^, with more substantial tails on the left, indicating that the upper limit of bulge is “harder” than its lower limit: high-bulge/low-span (tighter) turns are more likely to be associated with conformational strain and steric clashes within the turn than low bulge/high span turns. **(c)** Skew: types {I, I’, VIb} and {II, II’, VIa1} can be formed into two loose groups that peak near skew 0 and .1 respectively, while types VIII and VIa2 peak to the left and right of these groups, and the “catch-all” type IV covers the range of all structurally classified types. **(d)** The warp distributions form five groups from left to right, consisting of types {II, II’}, {IV, VIII}, VIb alone, {VIa1, VIa2}, and {I, I’}. Groups {II, II’} and {IV, VIII} exhibit positive Pearson skewness, with long tails to the right, while {I, I’} shows negative skewness. The general similarity between the type IV and VIII distributions of span, warp and particularly bulge is notable.

### 3.3 Ramachandran-space parameter heatmaps

The distributions of geometric parameter values across the Ramachandran spaces of the backbone dihedral angles of the two central turn residues reveal the bond rotations that drive parameter variation. Figure 3 displays six examples of heatmaps of these distributions in the classical turn types or BB clusters; see the figure caption for interpretations. The ExploreTurns tool gives access to Ramachandran-space heatmaps for all parameters across all types, BB clusters and the global turn set.

**Figure 3.**
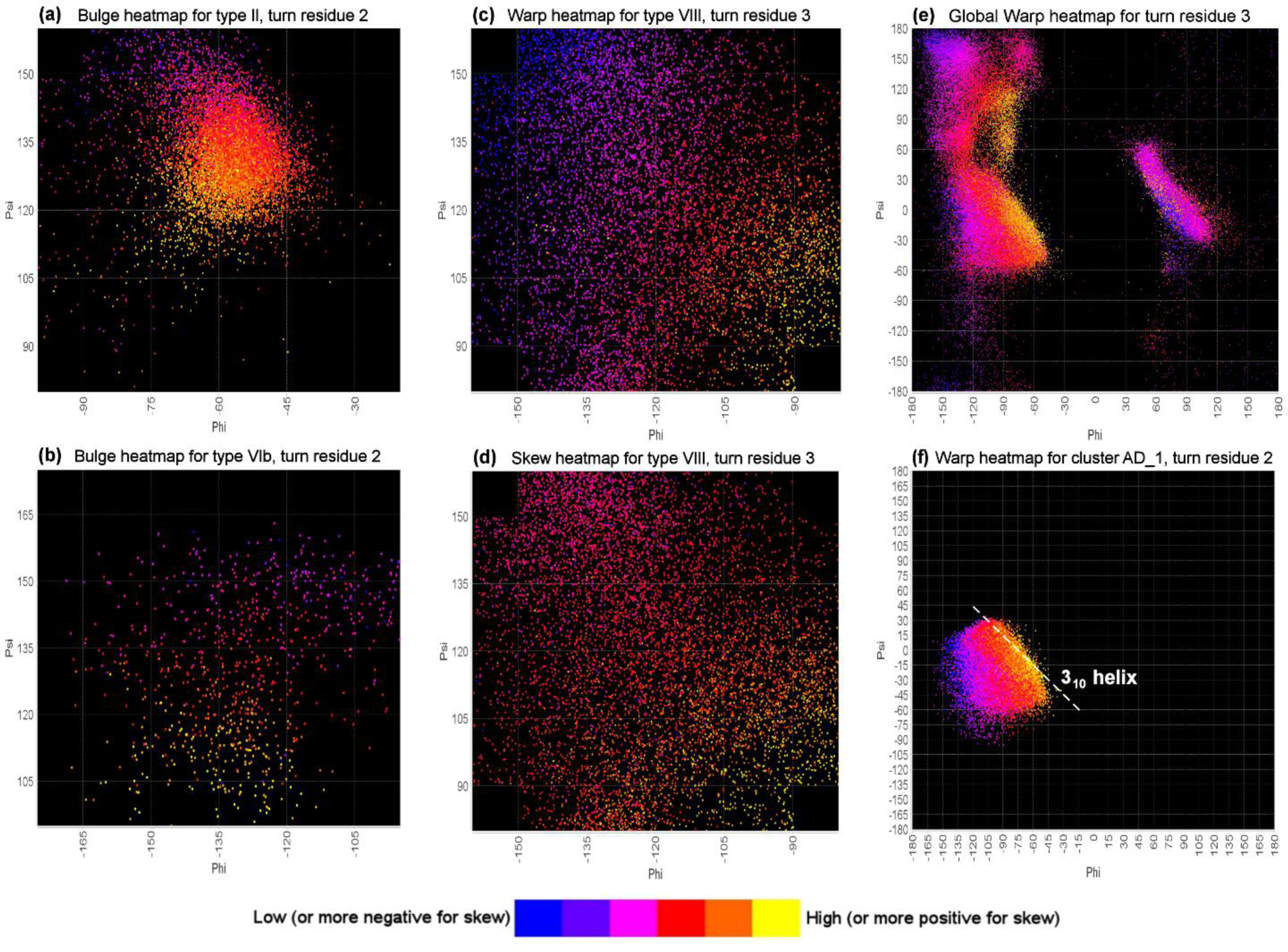
Ramachandran-space heatmaps reveal the bond rotations that drive parameter variation. Parameter values are colored on relative scales within each plot according to the legend; actual values can be obtained from ExploreTurns. **(a)** At the second residue of type II turns, rising Ψ_2_reduces bulge by rotating the *C*_α2_→*N*_2_ bond “outwards and upwards” (towards-y, +z), effectively reducing the x components of the three main-chain-path bonds in the plane of the first peptide bond. **(b)** The second residue of type VIb turns shows a more uniform gradient of falling bulge as Ψ_2_ rises, driven by an almost directly outwards rotation of *C*_α2_→*N*_2_. **(c, d)** The warp and skew heatmaps for residue 3 in type VIII turns show broad, highly correlated gradients of falling warp and increasingly negative skew from lower-right to upper-left as the residue opens into a more extended conformation, both flattening the turn and increasing its C-terminal half-span (and consequently its N-terminal “lean”). **(e)** The most characteristic feature of high-warp turns is the orientation of the second peptide bond approximately parallel to the z axis, and the global warp distribution at residue 3 shows a falling gradient over a wide range of Ψ_3_ as φ_3_ becomes increasingly obtuse below −60°, accompanying the rotation of the peptide bond down towards the turn plane. **(f)** In the type I-associated BB cluster AD_1, which contains about half of all turns, warp rises as φ_3_ becomes increasingly positive (more acute), supporting the sharper BB reversals required at *C*_α3_to direct the BB back down towards the turn plane after the large z-dimensional excursions associated with high warp. Warp is maximal near the line along which (φ, Ψ) sums to −75°, ideal for a 3_10_ helix, demonstrating the relationship between warp and helical character; a 3_10_helix can be viewed as an extension of a (high-warp) beta turn^19^.

### 3.4 H-bond energy vs. parameter value

Figure 4 plots the electrostatic (dipole-dipole) energy of the 4>1 BB H-bond versus geometric parameter values for classical types I and II, the largest types in which the H-bond is common. In type I, energy drops steadily with increasing warp, from .3 kcal/mol for the flattest turns to −1.8 kcal/mol at high warp (Figure 4a), while in type II energy shows a gradual increase as warp rises (Figure 4b). The negative correlation (*r* = −.61) between H-bond energy and warp in type I can be rationalized by the observation that the 4>1 H-bond donor and acceptor typically show a much poorer alignment, with a greater H…0 distance, in flatter vs. higher-warp turns due to rotations of the first and third peptide bond planes as warp falls; the primary factor is likely the rotation of the *C*_1_→0_1_ H-bond acceptor towards-z, but the *N*_4_→*H* donor can also exhibit a limited rotation towards +z. The incompatibility between low warp and the 4>1 H-bond in type I turns is a major contributor to the much lower frequency of this H-bond in type I (71%) than in the other three largest classical types that commonly exhibit it: I’ (95%), II (86%), and II’ (92%); H-bond frequency in the 47% of type I turns with warp greater than .75 is 90%.

**Figure 4.**
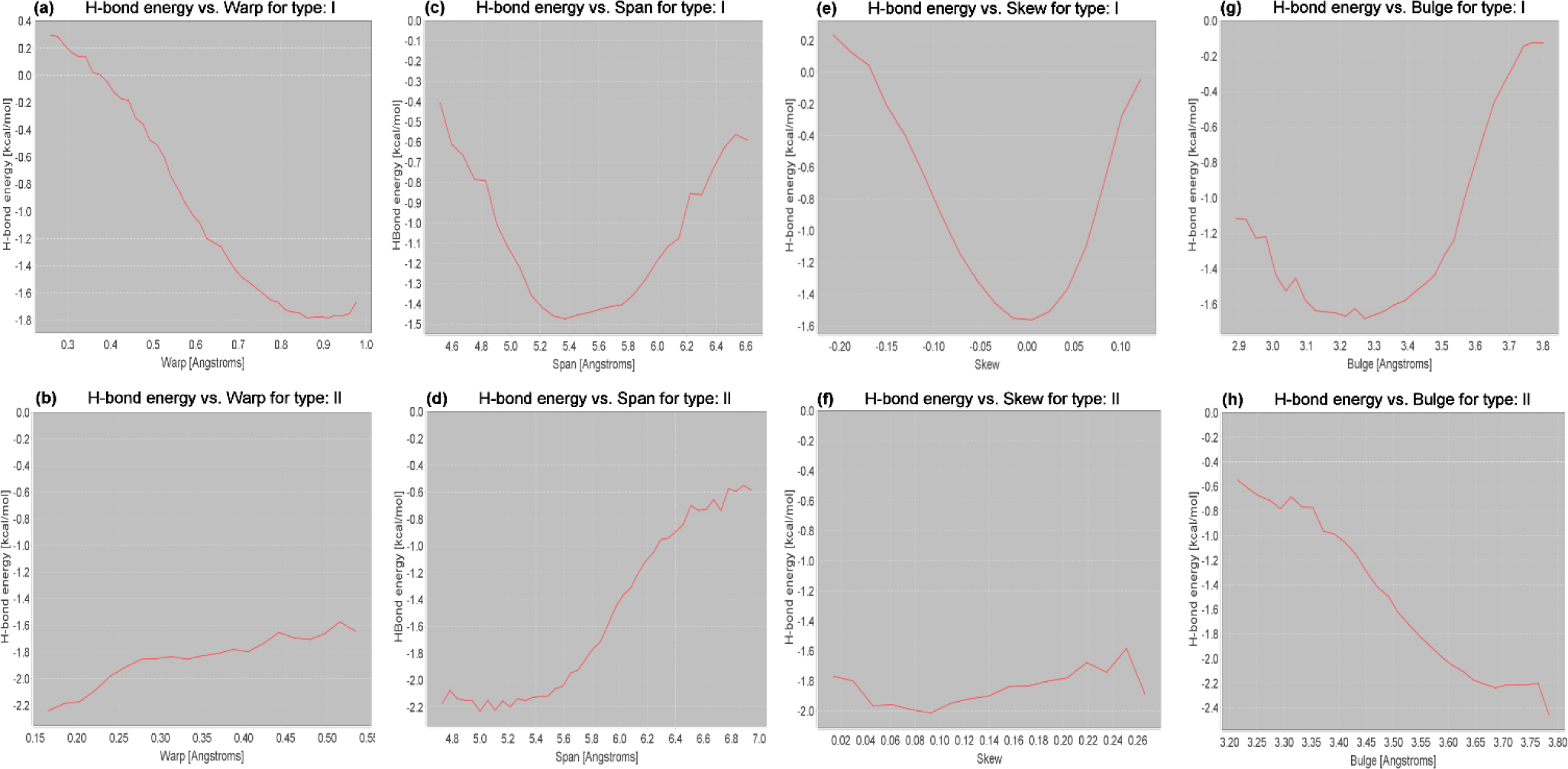
Electrostatic (dipole-dipole) energy of the 4>1 beta-turn H-bond vs. parameter value for span, bulge, skew and warp in classical types I and II. **(a, b)** Energy rises steeply with falling warp in type I, reflecting conformational changes that disrupt the H-bond as the turn flattens. By contrast, energy falls gradually as warp drops in type II. **(c, d)** Energy rises as span rises above 5.4Å in both types, reflecting H-bond lengthening, but in type I energy also rises as span falls below 5.3Å, reflecting an association between low span and (H-bond disrupting) low warp. **(e, f)** Energy shows only a weak dependence on skew in type II, but rises strongly with absolute skew in type I. **(g, h)** Energy falls steadily with increasing bulge in type II, in agreement with a simple wire-frame model in which span falls as bulge rises, but in type I energy increases as bulge climbs above 3.3Å, reflecting an H-bond-disrupting drop in warp with increasing bulge.

In type I (Figure 4c), energy falls as span increases until a minimum is reached near span 5.4Å, but then reverses its trend and rises steadily, while in type II energy remains at its minimum as span increases until about 5.4Å, then rises steadily (Figure 4d). The rise in H-bond energy above 5.4Å in both types can be rationalized by the expected increase in H-bond length with increasing span, but the initial energy drop with rising span in type I defies this; it is likely best explained by the correlation between span and warp in this type (Supplementary Section 2.2): lower span is associated with lower warp, which disrupts the H-bond.

H-bond energy is only weakly sensitive to skew in type II (Figure 4f), with low energy across the skew range, but in type I (Figure 4e) energy has its minumum at skew zero and rises dramatically with absolute skew.

The steady drop in H-bond energy in type II as bulge increases (Figure 4h) can be explained as an indirect effect, operating through the negative correlation between bulge and span, that is consistent with the behavior of a simple wire-frame model for the turn: a larger bulge results in a smaller span, which shortens and strengthens the H-bond. However, this picture does not hold in type I (Figure 4g), where energy falls to a minimum near 3.2Å, then begins a steady rise from 3.3Å upwards. This behavior can also be rationalized as an indirect effect, this time operating through a strong negative correlation between bulge and warp in this type (*r* = −.77): rising bulge, like falling span, is associated with falling warp, which disrupts the H-bond.

### 3.5 Parameter dependence of sequence motifs

The overrepresentations of multiple key sequence motifs in beta turns and their BB neighborhoods show clear dependencies on parameter values. Figures 5, 6 and 7 present plots of fractional overrepresentation vs. span, bulge, skew and warp for sequence motifs which commonly form SC/BB H-bonds (Figures 5, 6), SC/SC H-bonds (Figure 7a,b), or pi-stacking interactions (Figure 7c); see the figure captions for discussion. As the plots show, turn parameters can serve as structural discriminators within classical types or BB clusters that dramatically increase the specificity of measurements of sequence preferences in turns and their BB neighborhoods. For example, while the sequence motif that specifies Thr at turn position 3 (T3) is overrepresented by 48% in the set of all type I turns, Figure 6e shows that its overrepresentation peaks near 320% at low warp, reflecting roles that include SC/BB H-bonds linking the flat (for type I) “chair back” turn in the type 2 beta-bulge loop (BBL2)^20^ to the motif’s bulge. Similarly, the pair motif D1T3, associated with SC/SC H-bonding above the turn plane, is overrepresented by just 9% in the set of all type I turns, but Figure 7b shows that it peaks near 100% at high warp, which is likely favorable because it raises Thr’s SC into position for H-bonding with Asp.

**Figure 5.**
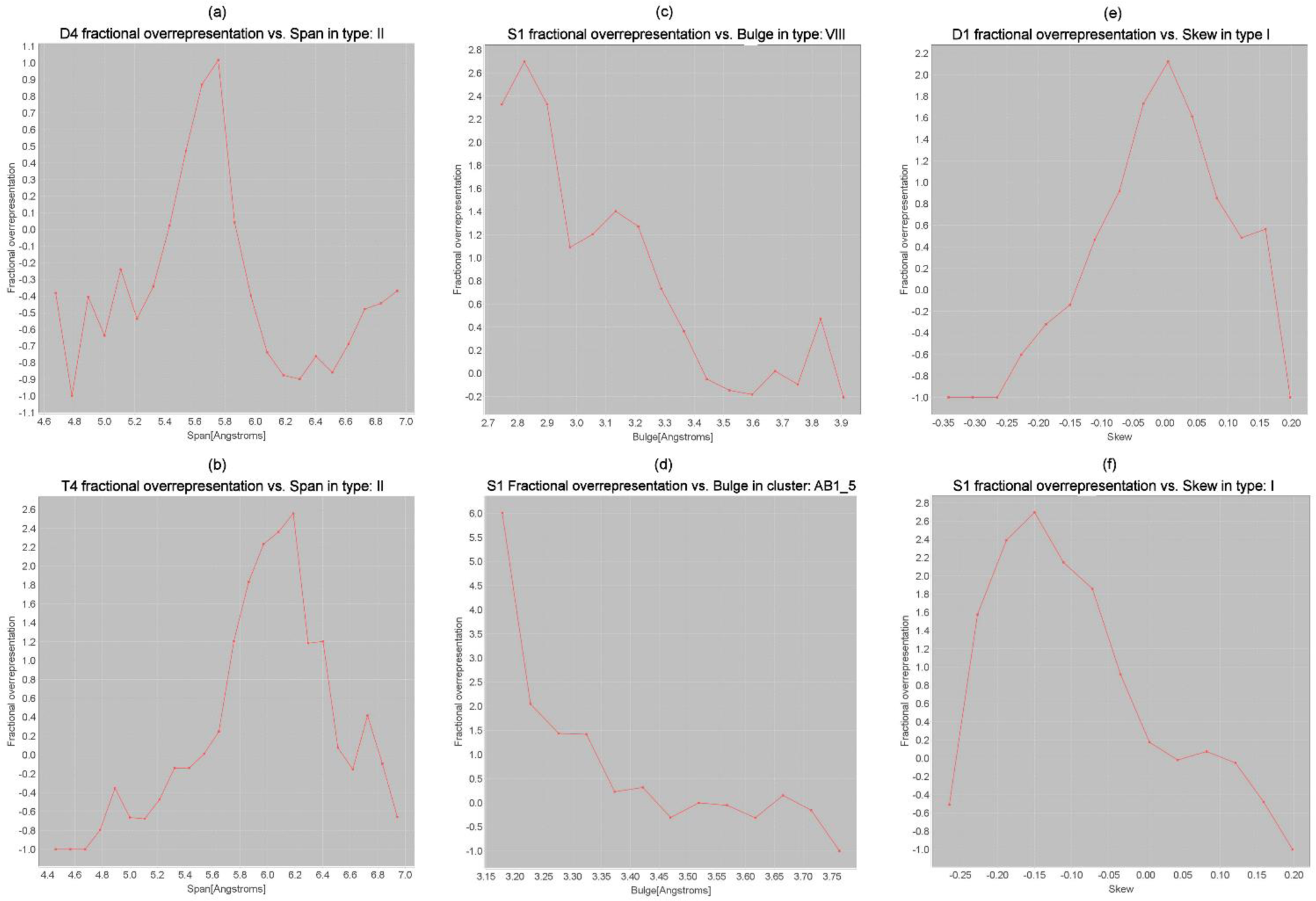
Examples of span, bulge and skew dependence of SC motifs. **(a, b)** Plots of the fractional overrepresentations of Asp and Thr at turn position 4 (D4 and T4) vs. span in type II turns show prominent maxima that likely correspond to the optimal spans for SC/BB H-bonding with the main-chain NH group of the first turn residue (MCN1, for D4) or the main-chain carbonyl group of that residue (MCO1, for T4). **(c)** The overrepresentation of Ser at position 1 (S1) drops below zero as bulge rises in type VIII, likely reflecting the lengthening and weakening of the H-bond between Ser’s SC and MCN3 (adjacent to the turn center) in this ST turn/motif^21^. **(d)** S1’s overrepresentation also falls as bulge increases in the BB cluster labelled here as AB1_5, which is one of the four clusters representing classical type VIII in the cluster classification (and the fifth largest cluster in the ultra-high resolution dataset used to derive the system^4^). **(e, f)** While Asp at position 1 (D1) in type I turns shows peak overrepresentation near zero skew, Ser at position 1 (S1) shows a negative-skew bias; this contrast may reflect the suitability of a short N-terminal half-span (which promotes negative skew) for optimal H-bonding between Ser’s shorter SC and the BB NH group at the turn’s C-terminus (MCN4).

**Figure 6.**
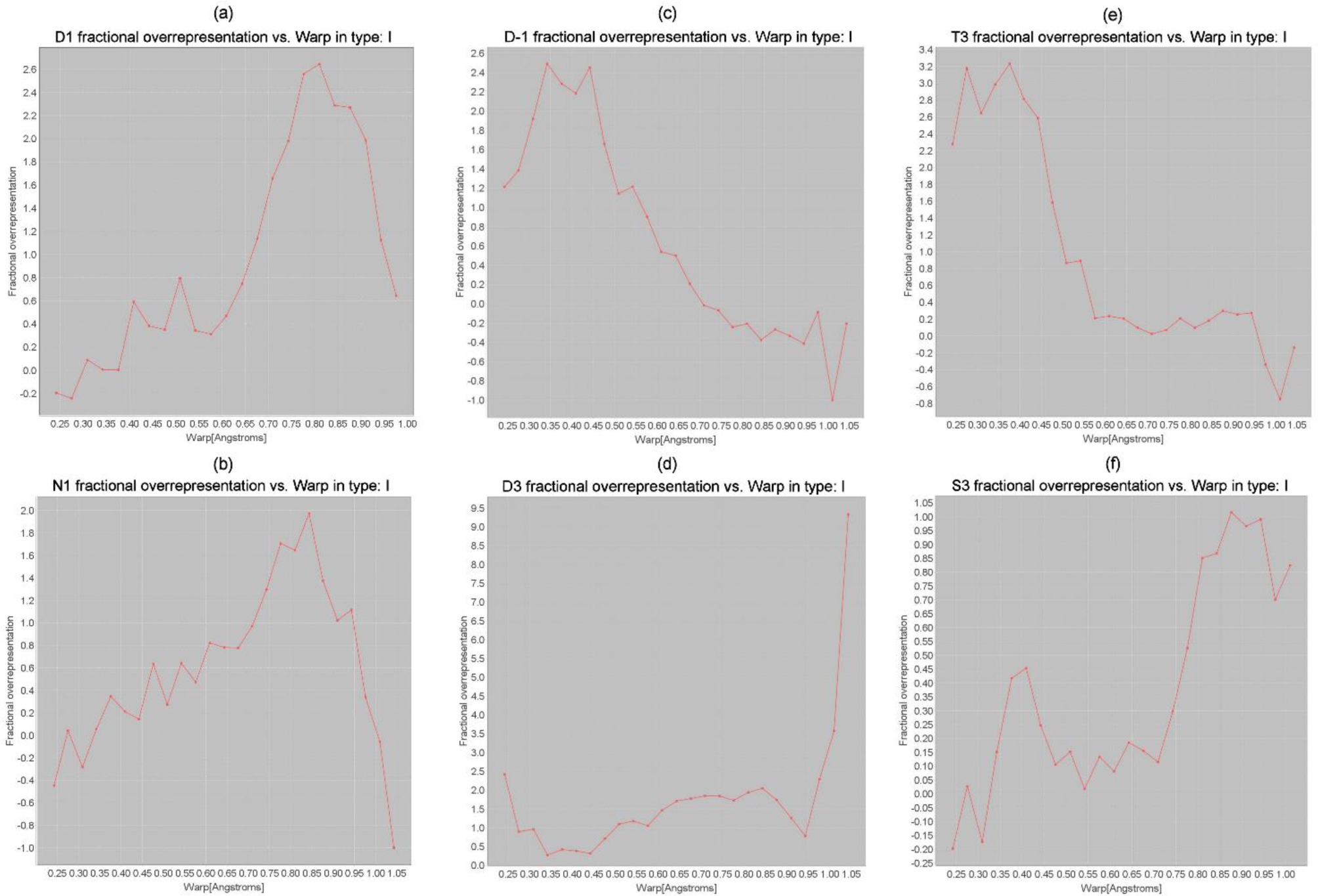
Examples of warp dependence of SC motifs. **(a, b)** Asp and Asn at turn position 1 (D1 and N1) in type I turns show peak overrepresentation at high warp, reflecting the compatibility of this geometry with SC/BB H-bonding with MCN3/MCN4 in Asx turns/motifs^22^ in a multiple contexts including open loops and type 1 beta-bulge loops^20^ (BBL1) (see Figure 9g). The favorability of high warp for these motifs is likely due, at least in part, to the strong negative correlation between warp and bulge (Supplementary Section 2.2), which results in a shorter distance between these SCs and MCN3 at high warp. **(c)** In contrast to D1/N1, Asp just before the turn (D-1) peaks at low warp, reflecting compatibility with structures such as the type 2 beta-bulge loop (BBL2, Figure 9h) and the alpha-beta loop^23^, in which flatter turns orient at approximately 90° to the incoming BB; D-1’s SC supports these “upright turns” by H-bonding with MCN2/MCN3. **(d)** The plot for Asp at turn position 3 (D3) reflects multiple roles: at minimum warp, the motif links the upright turns in BBL2s to MCN+1 in the loop’s bulge, at mid/high warp D3 H-bonds with its own BB NH group and/or MCN+1 in BBL1s and other structures, while at maximum warp the motif H-bonds with MCN+1 and the SC immediately after the turn in the BBL1s of beta propellers^24^. **(e)** At low warp, Thr at position 3 (T3) forms SC/BB and SC/SC H-bonds in BBL2s; this motif is four times as common as D3 in this context (see Figure 9h). **(f)** Ser at turn position 3 (S3) peaks at low warp, reflecting a role like that of D3/T3 in BBL2s, and at high warp, reflecting roles including SC/SC H-bonding with turn position 1 in multiple contexts and SC/BB H-bonding with MCN+1 in structures such as BBL1s.

**Figure 7.**
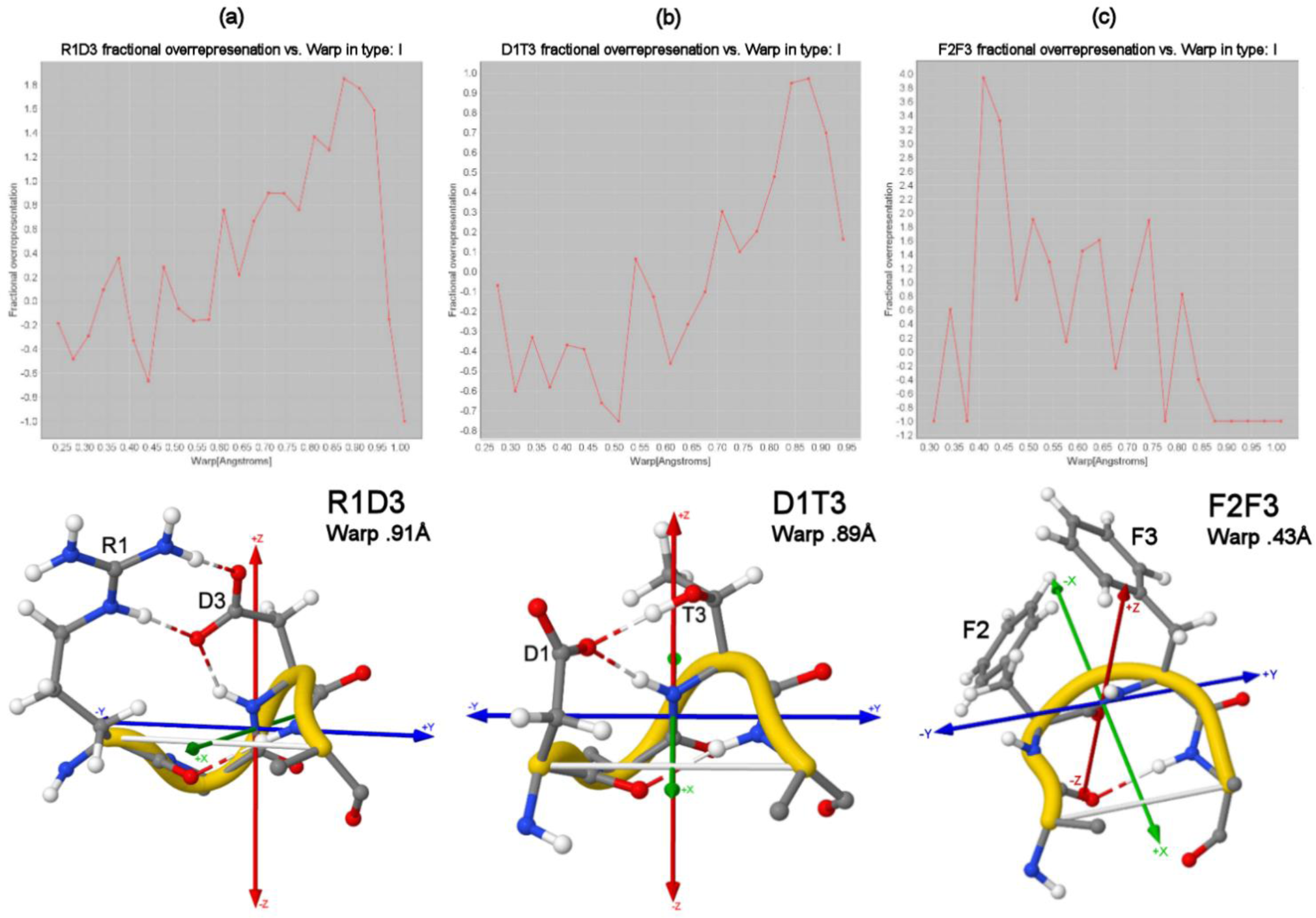
Examples of warp dependence of pair SC motifs. **(a, b)** Arg1+Asp3 (R1D3) and Asp1+Thr3 (D1T3) show strong preferences for high warp, which likely optimizes SC positioning for H-bonding (4A56_A_55 and 3AL2_A_1301). **(c)** Phe2+Phe3 (F2F3) favors lower warp, which brings the SCs closer for perpendicular pi-stacking (3PYC_A_29).

With the guidance of built-in motif overrepresentation plots like those shown here, turn parameters can be used in ExploreTurns to tune structures for compatibility with particular sequence motifs, and the facility’s motif detection tools can be used to identify the motifs important in particular types, clusters or BB contexts. Sequence motifs can be comprehensively explored using the interactive conformational heatmaps generated by the MapTurns tool; to open a motif’s map, enter its label into the *Sequence motif* box in ExploreTurns and click ***Map Motif***.

### 3.6 Effect of *cis*-/cis-Pro peptide bonds on bulge and skew

Figure 8 demonstrates strong relationships between the occurrence of *cis-/cis-Pro* peptide bonds in a beta turn and the turn’s skew and bulge, and these effects show a degree of superposition when *cis* conformations are combined (see the figure caption for details).

**Figure 8.**
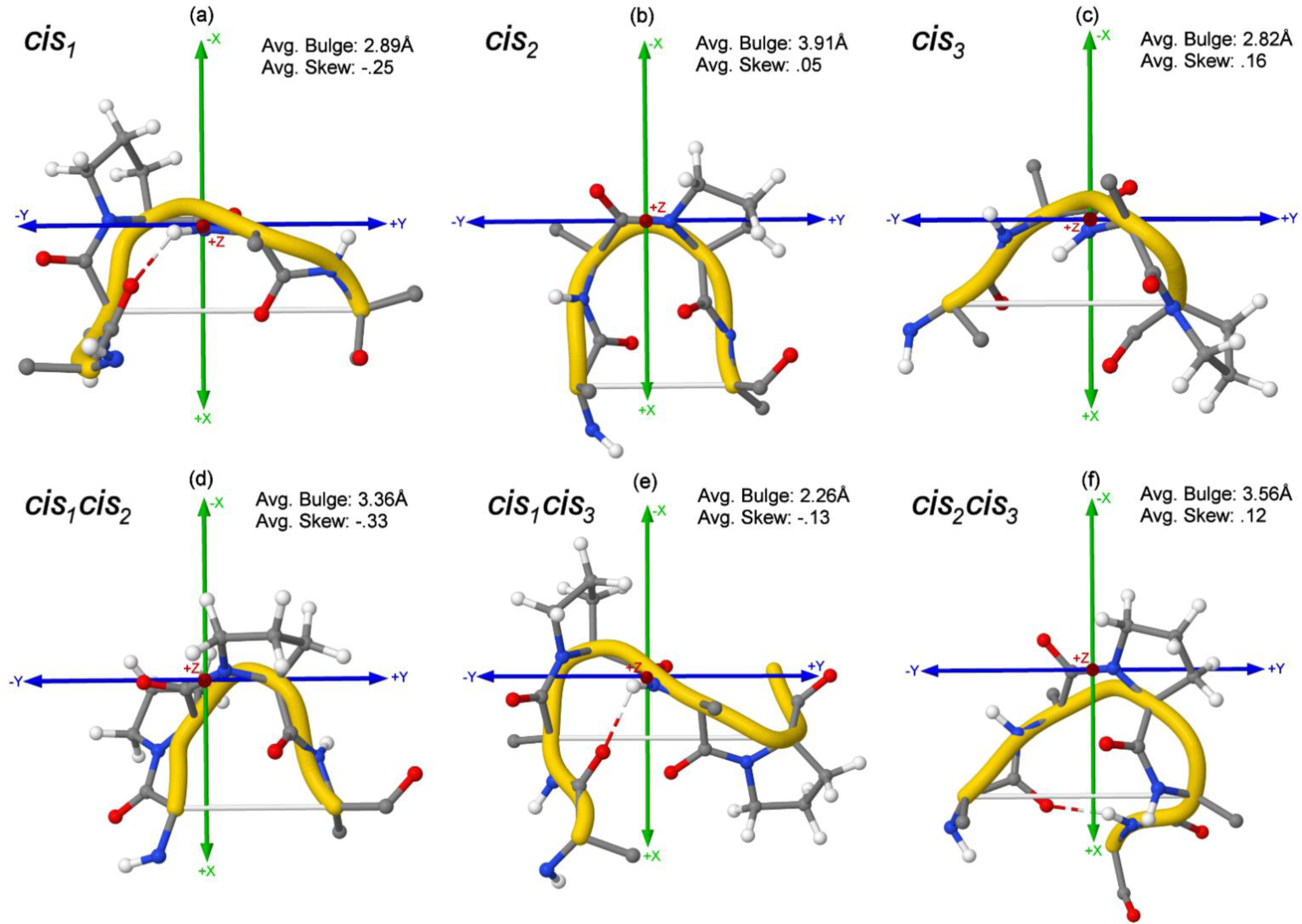
*cis*-/cis-Pro-peptide bonds influence bulge and skew and show a degree of superposition. **(a)** Turns with a *cis* conformation at the first peptide bond (*cis*_1_) exhibit low average bulge (2.89Å vs. the 3.45Å global avg.) and a negative-skew bias (3O4P_A_140). **(b)** *cis*_2_ turns show high bulge with a minor positive-skew bias (1SG6_A_99). **(c)** *cis*_3_ turns show low bulge and a positive skew bias (3UCP_A_685). **(d, f)** In *cis*_1_*cis*_2_ and *cis*_2_*cis*_3_ turns, the low-bulge biases of the component *cis*_1_ and *cis*_3_ peptide bonds effectively cancel the high-bulge bias of *cis*_2_, yielding bulges close to the global mean, while skew biases of the same signs as the *cis*_1_ and *cis*_3_ components persist (enhanced in *cis*_1_*cis*_2_) when combined with *cis*_2_’s near-neutral skew (3LCC_A_180, 4OW5_A_102). **(e)** In *cis*_1_*cis*_3_ turns, the low-bulge biases of the *cis* components show an additive effect, resulting in extreme low bulge, while *cis*_1_’s larger negative-skew bias is reduced by combination with *cis*_3_’s smaller positive bias (1ZR6_A_397). Sample sizes are small for all combination *cis*-types. Note that for all types except *cis*_2_*cis*_3_, a proline residue follows the *cis*-peptide bonds in a majority of structures, and the results shown here combine the effects of Pro and non-Pro *cis*-peptide bonds. However, analysis with ExploreTurns shows that individual non-Pro (*cis*_1_, *cis*_2_, *cis*_3_)-peptide bonds show relationships to bulge and skew similar to the combined results shown in the top row of the figure as well as the results seen for individual Pro *cis*-peptide bonds.

The skew biases of *cis*-peptide bonds can be compounded by combination with biased Ramachandran types. For example, the combination of a *cis* conformation at the first peptide bond (*cis1*, avg. skew −.25) with the negatively-biased type AB (-.18), yields *cis*AB (-.41), and the combination of *cis*3 (.16) with Ba (.20) yields Ba*cis* (.36), in which the biases add almost exactly. The Ramachandran type that specifies *cis*-polyPro conformations at both central turn positions (*cis*P*cis*P) shows extreme skew bias (-.46).

To view a Ramachandran-region map in ExploreTurns, click ***Ramachandran Types***; turns of a particular Ramachandran type can be browsed by entering the type name in the ***Type or BB cluster*** box.

### 3.7 Parameter correlations

The structural constraints imposed by bond lengths and angles in the beta-turn BB impose relationships between the parameters that reflect geometric tradeoffs in turn design. Supplementary Section 2.2 presents a table of parameter cross-correlations and rationalizes several key examples.

### 3.8 Roles and contexts of parameter variation

Turn parameters describe geometric variations which effect the suitability of turns for particular roles and contexts. The highly anti-correlated bulge and span parameters are major determinants of a turn’s curvature, and turns of high bulge/low span are found where the BB must reverse within spaces that are narrow in the y dimension of turn coordinates. Bulge provides variability in the x dimension; turns with higher bulge can project the BB and SC groups of a turn’s central residues further toward -x to support structural interactions, participate in ligand binding or active sites, or increase solvent exposure. On the opposite end of the bulge/span scale, the most common context for turns with the largest spans is extended loops.

Skew is a measure of a turn’s asymmetry in the y dimension, while the half-spans that determine skew represent degrees of freedom in this dimension that provide some flexibility in the positioning of BB and SC groups on each side of the turn axis, which can be used to satisfy packing constraints or support interactions. Ramachandran types which specify a more extended conformation at one central turn residue are associated with skew biases, since they lengthen one half-span. For example, Ramachandran types AB, AZ and AP show negative-skew biases due to their {B, Z, P} components; types AB and AZ are largely responsible for the negative skew bias of classical type VIII. Extended conformations at the second turn residue have the opposite effect on skew: types Ba, Za, Pa and Pd show positive-skew biases, and Pa and Pd are responsible for most of classical type II’s positive bias.

Warp measures the magnitude of the excursions a turn’s BB makes above and below the turn plane; low warp can accommodate tight packing in the z dimension, while high warp enables a turn’s BB and SC groups to interact with structures farther above and/or below the plane.

Just as the values of all four parameters show particular regimes that are compatible with key SC motifs, they also exhibit compatibilities with the structures in a protein that incorporate turns, including beta sheets, local H-bonded motifs^23^ and ligand-binding sites. Turn geometry and context are interrelated; a turn’s AA content can bias its conformation in support of a context, while the H-bonding, backbone stress or ligand binding associated with a context can influence the turn’s geometry, affecting both parameter values and turn type. Figure 9 displays examples of structural contexts associated with higher or lower parameter values.

**Figure 9.**
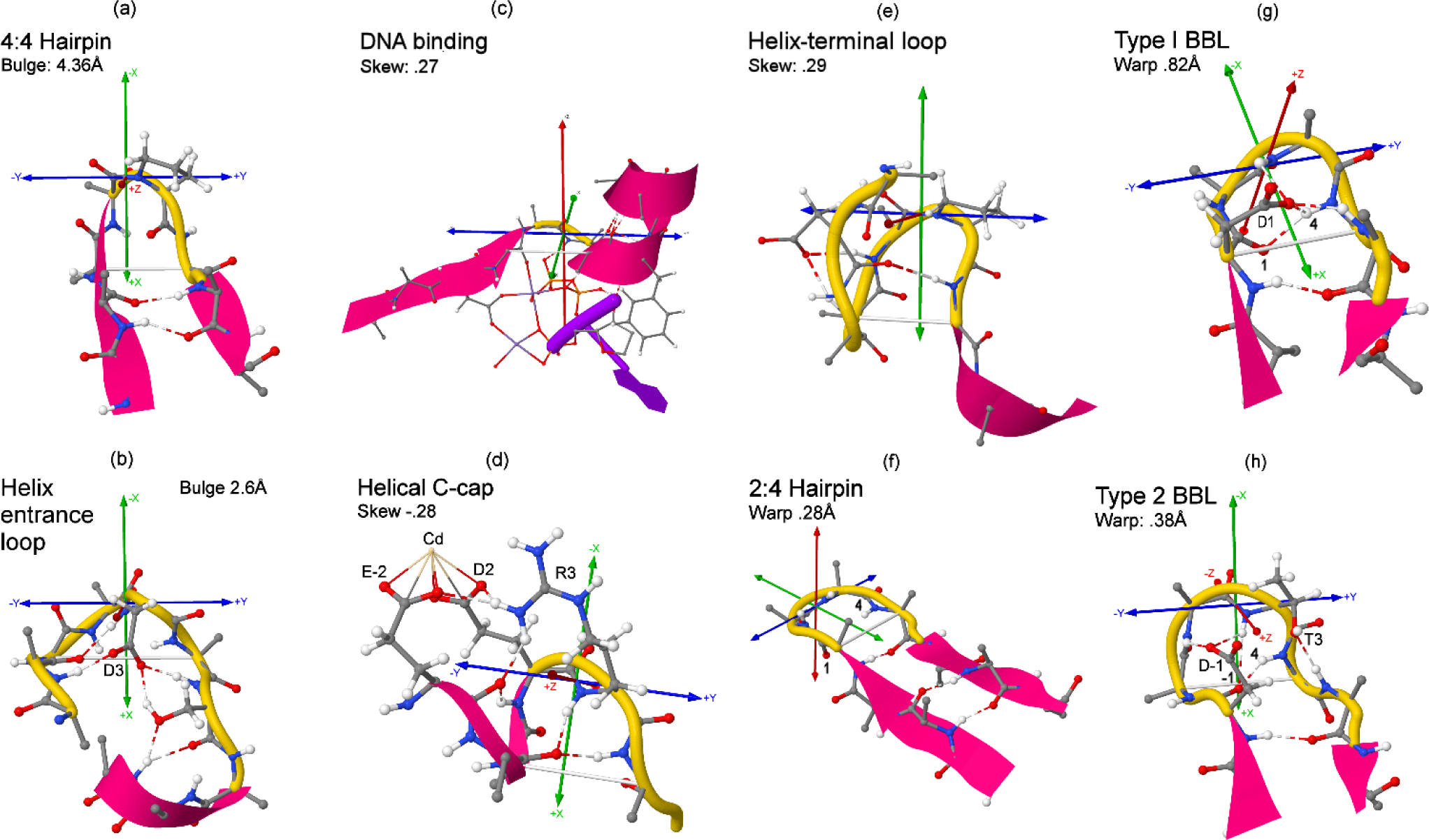
Roles and contexts of parameter variation. **(a)** Chain-reversing loops in 4:4 beta hairpins are commonly formed of high-bulge type VIb (mostly Ramachandran type B*cis*P) turns (3AKS_A_81). **(b)** The low bulge of a type IV (dA) turn in a helix-entrance loop positions the Asp SC at turn position 3 (D3) to mediate a network of H-bonds binding the loop (5UJ6_A_542). **(c)** The positive-skew geometry of a type IV (Ba) turn, with its shorter C-terminal half-span, helps position turn and helix groups for DTP/DNA binding in DNA polymerase (5KFZ_A_14). **(d)** The shorter N-terminal half-span of a negatively-skewed turn helps position the helix with respect to the turn for C-capping and Cd binding (3B40_A_50). **(e)** The short C-terminal half-span of a positively-skewed type VIa2 (B*cis*A) turn (avg. skew .25 for the type) shortens a BB H-bond that secures the geometry of a helix-terminal loop (3V5C_A_287). **(f)** The low warp of a type I turn simultaneously enables compatibility with the ladder component of a hairpin and disrupts the turn’s 4>1 H-bond, yielding a 2:4 hairpin in place of the more common 2:2 type. **(g)** A high-warp type I turn is compatible with the BBL1’s interior 4>1 BB H-bond as it forms the back of the loop’s “reclined chair”; the motif is shown here with Asp at position 1 (D1, see Figure 6a) (4K9Z_A_34). **(h)** A low-warp type I turn is compatible with the “upright chair” conformation of a BBL2 and the motif’s characteristic 4>-1 H-bond. The loop is shown with its chair-back turn supported by SC/BB H-bonds mediated by Asp just before the turn and Thr at turn position 3 (D-1 and T3, see Figure 6c,e) (3SCY_A_360).

The 2:4 beta-hairpin^25^ (Figure 9f) demonstrates a notable relationship between warp and beta-turn context. Most chain-reversing turns in 2:2 hairpins (the most common hairpin type) are of types I’ or II’; turns of type I are much less common due to a geometric clash between the right-handed inter-strand twist of the hairpin’s beta ladder and the left-handed twist between the turn’s incoming and outgoing BB segments^7^. Low warp relieves this clash by reducing the turn’s twist, rendering its structure more compatible with the ladder, but the conformational changes associated with warp reduction commonly disrupt the turn’s 4>1 H-bond (Figure 4a), yielding a 2:4 hairpin in place of the more common 2:2 type. As a result, more than half of 2:4 hairpins in the dataset in effect owe their existence to the compatibility between low warp and the ladder’s twist.

The chain-reversing turns in type 1 and 2 BBLs, which are both commonly of type I, demonstrate further relationships between warp and context. In the BBL1 (Figure 9g), a high-warp turn supports the loop’s 4>1 H-bond, commonly forming a “reclined chair”, while in the BBL2 (Figure 9h), the low-warp geometry of the turn that forms the back of the motif’s “upright chair” suppresses the 4>1 H-bond in favor of the 4>-1 alternative, which is characteristic of the loop.

### 3.9 Parameters and structure quality

Since geometric parameters enhance the description of turn geometry in Euclidean space, they may aid the detection and characterization of structurally suspicious conformations. Supplementary Section 2.3 investigates the relationship between parameter values and structure quality, and finds associations between extreme values and an increased frequency of geometric quality criteria with outliers as measured by PDB validation tools^26^.

## 4 Conclusions

The turn-local coordinate system and global alignment derived here resolve the difficulties with visualization inherent in the Ramachandran-space turn representation, providing a natural framework for the comparison and analysis of the wide variety of beta-turn structures and the contexts in which they occur, including H-bonded motifs, ligand-binding/active sites and supersecondary structures. The utility of this framework has been demonstrated by the application of ExploreTurns^6^ to the characterization of multiple new short H-bonded loops, the mapping of sequence preferences in Asx N-caps, and the profiling of Schellman loop/beta turn combinations, and the framework also supports the comprehensive maps of sequence motifs produced by MapTurns^14^.

Geometric parameters provide a Euclidean-space description of the bulk BB shapes of turns that reflects meaningful modes of geometric variation not explicitly captured by Ramachandran-space representations. Parameters complement existing turn classification systems, enabling a fine-tuning of backbone geometry within classical types or BB clusters that supports the selection of structures for compatibility with particular SC interactions or turn contexts and produces major improvements in the specificity of measurements of AA sequence preferences in turns. Since close to two-thirds of residues in protein loops are found in beta turns^4^, parameters can bring needed additional order to the characterization of the irregular geometries of loops, which represent about half of all protein structure^27^.

Turn-local coordinates and geometric parameters should prove useful in any application where an enhanced Euclidean-space picture of turns and structural tuning can improve understanding or performance. One potential area of application is protein design, and in particular the design of binding and active sites, where turns are common and even small changes in geometry may have major impacts on function. Parameters may also prove valuable in structure validation, by improving the characterization of suspicious conformations and identifying extreme geometries that may not always be highlighted as outliers by existing validation tools.

## Declarations

### Availability of data and materials

All data supporting the conclusions of this article are available from the Protein Data Bank at www.rcsb.org. The ExploreTurns, MapTurns and ProfileTurn web tools are available at www.betaturn.com.

## Supporting information

Supplementary sections

## Acknowledgements

The author wishes to thank Athena Newell for useful discussions and work as a research assistant and Kit Newell for work as a research assistant. The constructive suggestions and support of the attendees of ISMB/3DSig 2019 (Basel) and the 33rd and 37^th^ annual symposia of The Protein Society (2019 Seattle and 2023 Boston) also contributed to the development of this project.

## Supplementary Material

**Supplementary file 1** (PDF). Supplementary sections, with table of contents.

## Abbreviations

AA: Amino acid
BB: Protein backbone
SC: Amino acid side-chain
Position index: Specifies residue positions in the turn and its four-residue N-/C-terminal BB neighborhoods according to the key: −4-3-2-1+1+2+3+4, where the brackets delineate the turn.
MCN + position index: Main-chain NH-group H-bond donor at the specified position. For example, MCN-1, MCN2, and MCN+3 refer to the BB H-bond donors one residue before the turn, at the second turn residue, and three residues after the turn respectively.
MCO + position index: Main-chain carbonyl oxygen H-bond receptor at the specified position.
BB H-bonds: BB H-bonds are specified by indicating the donor and acceptor positions in the turn frame, separated by ’>’. For example, “4>-1” specifies an H-bond donated by MCN4 and accepted by MCO-1.
AA sequence motif: AA sequence motifs are labelled with the single-letter codes of their included AAs, each followed by its position in the turn frame. For example, “D-1T3” specifies Asp just before the turn and Thr at the third turn position.
PDB address: PDB addresses for turn structures are given in the format ABCD_X_N, where ABCD is the 4-character PDB file name, X is the chain letter, and N is the position in the chain of the first turn residue.
BBL: Beta-bulge loop; also BBL1 (type 1), BBL2 (type 2).

