## Supplementary sections for "A geometric parameterization for beta turns"


1 | Methods 2
- 1.1 Accuracy of the global turn alignment 2
- 1.2 The beta-turn dataset 2
- 1.3 H-bond definition 2
- 1.4 Computing sequence motif overrepresentation 3

2 | Results 4
- 2.1 Parameter distributions in the BB clusters 4
  - Figure S1. Parameter distributions in the first nine BB clusters 5
  - Figure S2. Parameter distributions in the last nine BB clusters 6
- 2.2 Parameter correlations 6
  - 2.2.1 Cross-correlation matrices 6
    - Table S1. Cross-correlation matrices for the classical types 7
  - 2.2.2 Statistical significance of parameter correlations 8
- 2.3 Parameters and structure quality 9
  - Figure S3. PDB outlier elevation at global parameter extremes 10
  - Figure S4. PDB outlier elevation at type I/I' extremes 11
  - Figure S5. PDB outlier elevation at type II/II' extremes 12
  - Figure S6. PDB outlier elevation at type VIII extremes 13

3 | Discussion 14
- 3.1 Effect of turn BB geometry on the interpretation and utility of the warp parameter 14

References 15

### 1 | Methods

#### 1.1 Accuracy of the global turn alignment

The accuracy of the global turn alignment implicitly established by the turn-local coordinate system was tested by a comparison between implicit pairwise turn alignments and the best alignments generated by a perturbative search procedure. In this process, the average distance between the corresponding BB atoms of a randomly chosen, implicitly aligned turn pair was compared to the distance obtained for the best alignment generated by a perturbative search applied to the pair which seeks the minimum average interatomic distance by introducing small variations in the relative position and orientation of the two turns. This procedure was applied to a random sample of 100 million turn pairs to generate an estimate of the average positional error of the implicit alignments. Since this estimate (.17Å) is comparable to the errors expected for the positions of well-determined atoms in well-refined structures (.1Å - .2Å; overall RMSDs for independently-determined 2Å structures have been measured as .5Å - .8Å)[1], the accuracy of the implicitly established alignment was judged to be satisfactory.

#### 1.2 The beta-turn dataset

Beta turns are defined here as four-residue BB segments with a distance of no more than 7Å between the alpha carbons of the first and fourth residues and central residues with DSSP[2] codes outside the set {H, G, I, E} that specifies helix or strand[3]. Turn/tail structures were extracted from their PDB files[4] with the aid of a list of peptide chains with a maximum mutual identity of 25% obtained from PISCES[5]. Since sequence motif detection identified multiple motif "artifacts" generated by remaining local redundancy in the data, an additional step of turn-local redundancy screening was applied: structures were clustered and filtered to reduce redundancy within five residues of turns to 40% or less; this threshold was chosen because it effectively reduced artifacts while preserving a dataset of sufficient size.

Turn structures were screened for quality by requiring that there be no missing residues within 5 residues of any turn and excluding structures with bond lengths that constituted outliers from established values. The final dataset of 102,192 turns was compiled with resolution and R-value cutoffs of 2.0Å and .25 respectively; average values in the dataset are 1.6Å and .20.

#### 1.3 H-bond definition

H-bonds are defined by both energy and geometric criteria. The electrostatic (dipole-dipole) energy of a BB H-bond is computed as[2]:

$$E = 27.888\left(\frac{1}{r_{ON}} + \frac{1}{r_{CH}} - \frac{1}{r_{OH}} - \frac{1}{r_{CN}}\right)$$

where the $r's$ represent the distances (Å) between the pairs of atoms indicated in the subscripts, and $E$ is expressed in kcal/mol. The DSSP energy cutoff of -.5 kcal/mol is applied.

Geometric H-bond criteria include a maximum H…O distance of 2.6Å, a minimum N-H…O (donor) angle of 100°, and a minimum H…O=C (acceptor) angle of 90°.

The measurement of H-bond energy and geometric criteria depends on the presence of amide hydrogens in the structures, which are added to the PDB files using Reduce[6]. Coordinates of hydrogen atoms (and all other atomic coordinates) are extracted for processing with Biopython[7]. For about .5% of residues, BB amide hydrogen atomic coordinates are not available, preventing the evaluation of H-bonds.

### 1.4 Computing sequence motif overrepresentation

Statistical models are used to evaluate the fractional overrepresentation, $(O - E)/E$, of single-AA and pair sequence motifs in sets of structures, where $O$ is a motif's observed count in the set and E is its expected count under a suitable null model.

For a single-AA sequence motif, the null model specifies that the probability $P$ of the motif's occurrence in the set is equal to the position-independent probability of the occurrence of the motif's AA anywhere in proteins, which is computed as the overall abundance fraction of the AA in the complete set of protein chains used in the study: $P = C_{AA}/T$, where $C_{AA}$ is the count of the AA in the set of all chains and $T$ is the total number of residues in the chain set. The motif's expected count in a structure set is then $E = NP$, where $N$ is the size of the set.

The null model for a pair sequence motif, which represents independence between the occurrences of the component AAs in the pair, specifies that the probability $P$ of the motif's occurrence in a structure set is equal to the product of the independent probabilities of the occurrences of its component AAs in the set: $P = (N_1/N) * (N_2/N)$, where $N_1$ and $N_2$ are the counts of the component AAs in the set and $N$ is the set size. The motif's expected count is $E = NP = (N_1 * N_2)/N$.

A larger fractional overrepresentation for a motif signals a lower likelihood that the null model of independence is correct, and a higher likelihood of a synergy between the motif's component AAs that represents an interaction such as an H-bond. P-values for sequence motifs are not computed here, but are available from the ExploreTurns tool (see the tool's user guide).

### 2 | Results

#### 2.1 Parameter distributions in the BB clusters

Figures S1 and S2 plot the parameter distributions in the eighteen BB clusters (which are ranked by size in the dataset used to derive the clusters[3]) together with the outlier category.

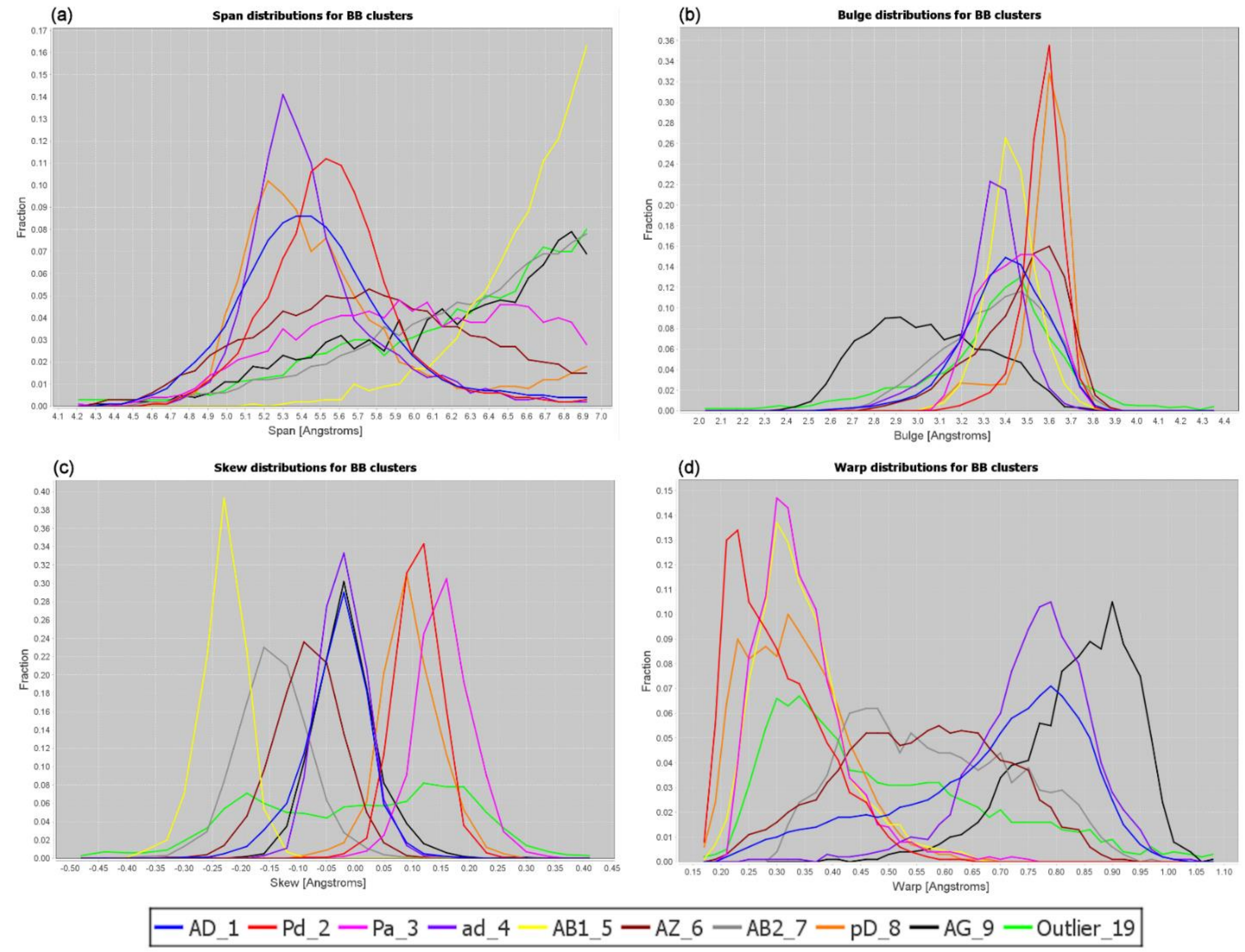


**Figure S1. Parameter distributions in the first nine BB clusters.** Parameter distributions in the first nine of eighteen BB clusters (ranked by size in the dataset used to derive the clusters[3]) and the outlier category (Outlier_19). **(a)** Span: of the clusters associated with classical type VIII (AB1_5, AZ_6, AB2_7, AG_9), clusters AB2_7 (grey) and AG_9 (black) show distributions most similar to that of type VIII (Figure 2a), climbing steadily to maxima at the maximum allowed span of 7Å, while the distribution of AB1_5 (yellow) climbs much more steeply, and AZ_6 (brown) contrasts sharply, with a broad distribution that peaks near 5.8Å. **(b)** Bulge: the distribution for the type-VIII associated cluster AG_9 (black) stands out, peaking well to the left of the other distributions. **(c)** Skew: the clusters that represent classical type II/II' (Pd_2, Pa_3 and pD_8) form a loose group on the right, while those that represent type VIII form a progression on the left covering a range of more negative skew, with AB1_5 (yellow) peaking at far left, AG_9 (black) in the center, near zero skew (overlaying with the type I/I' clusters AD_1 and ad_4), and the broader AB2_7 (grey) and AZ_6 (brown) falling between. **(d)** Warp: of the two clusters that represent classical type II, Pd_2 (red) peaks nearer to that type's maximum at far left, while Pa_3 (magenta) is offset towards higher warp. The very broad warp distribution of classical type VIII is partitioned into clusters AB1_5 (yellow), peaking at left near .3Å, AG_9 (black) peaking at far right near .9Å (the highest-warp peak for any type or cluster), and AZ_6 (brown) and AB2_7 (grey) peaking broadly in the center, spanning most of the plot.

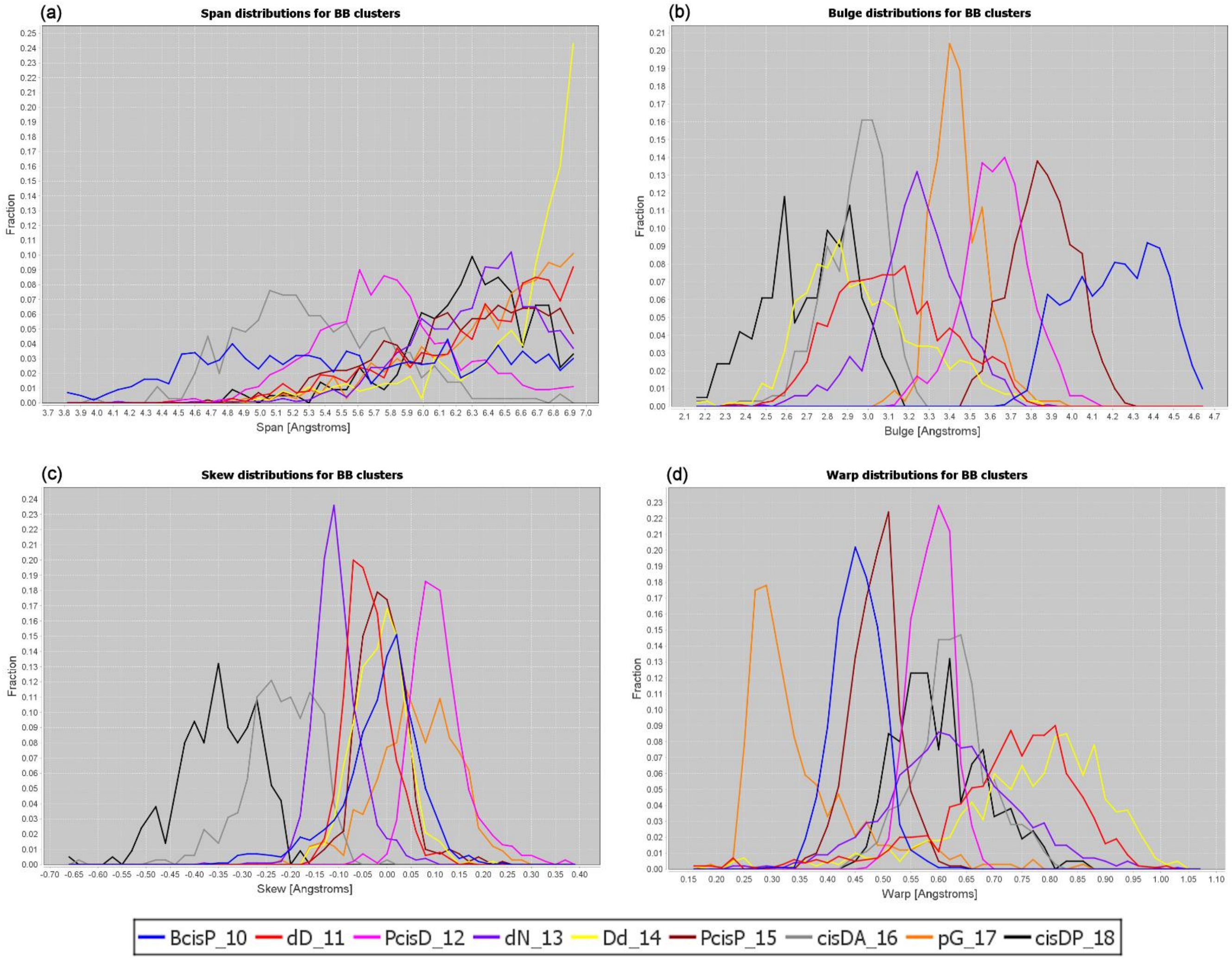


**Figure S2. Parameter distributions in the last nine BB clusters. (a)** Span: cluster Dd_14 (yellow) rises above all others as the 7Å limit of span is reached. **(b)** Bulge: the distributions form a progression covering a broad range of bulge. **(c)** Skew: clusters cisDP_18 (black) and cisDA_16 (grey) stand out at negative skew. **(d)** Warp: Cluster pG_17 (orange) stands out with a low-warp peak near .3Å, while the mirror-image clusters dD_11 (red) and Dd_14 (yellow) peak at high-warp near .8Å, and the other clusters cover a range at mid-warp.

### 2.2 Parameter correlations

#### 2.2.1 Cross-correlation matrices

The structural constraints imposed by the lengths and geometries of the bonds in the beta-turn BB enforce relationships between the turn parameters that reflect geometric tradeoffs in beta-turn design. Table S1 presents cross-correlation matrices of Pearson correlation coefficients ($r$) between turn parameters in all classical types, along with the corresponding values of $r^2$, which measure the fraction of the variance of one parameter in a pair that can be explained by a best-fit linear relationship with the other, and serve as a metric for the strength of the relationship.

| Type I | | | | |
|---|---|---|---|---|
| | Bulge | Warp | Skew | H-Bond Energy |
| Span | -.76/.58 | .33/.11 | .29/.09 | -.05/.00 |
| Bulge | - | -.78/.60 | -.33/.11 | .35/.12 |
| Warp | | - | .37/.13 | -.61/.37 |
| Skew | | | - | -.24/.06 |

| Type I' | | | | |
|---|---|---|---|---|
| | Bulge | Warp | Skew | H-Bond Energy |
| Span | -0.76/.57 | .23/.05 | .31/.10 | .22/.05 |
| Bulge | - | -.73/.54 | -.20/.04 | **.00/.00** |
| Warp | | - | **.04/.00** | -.22/05 |
| Skew | | | - | .09/.01 |

| Type II | | | | |
|---|---|---|---|---|
| | Bulge | Warp | Skew | H-Bond Energy |
| Span | -.88/.77 | .28/.08 | -.28/.08 | .57/.33 |
| Bulge | - | -.37/.14 | .21/.04 | -.63/.40 |
| Warp | | - | -.39/.15 | .25/.06 |
| Skew | | | - | .07/.00 |

| Type II' | | | | |
|---|---|---|---|---|
| | Bulge | Warp | Skew | H-Bond Energy |
| Span | -.89/.80 | .22/.05 | **-.07/.00** | .54/.30 |
| Bulge | - | -.35/.12 | **.03/.00** | -.56/.31 |
| Warp | | - | -.36/.13 | .14/.02 |
| Skew | | | - | .14/.02 |

| Type VIa1 | | | | |
|---|---|---|---|---|
| | Bulge | Warp | Skew | H-Bond Energy |
| Span | -0.87/.77 | .55/.30 | -0.34/.12 | .68/.46 |
| Bulge | - | -.64/.41 | .37/.13 | -.50/.25 |
| Warp | | - | -.30/.09 | .48/.23 |
| Skew | | | - | **.10/.01** |

| Type VIa2 | | | | |
|---|---|---|---|---|
| | Bulge | Warp | Skew | H-Bond Energy |
| Span | -.90/.81 | **-.03/.00** | -.55/.30 | -.74/.55 |
| Bulge | - | **-.14/.02** | .45/.20 | .74/.54 |
| Warp | | - | **-.03/.00** | **.21/.05** |
| Skew | | | - | .47/.22 |

| Type VIb | | | | |
|---|---|---|---|---|
| | Bulge | Warp | Skew | H-Bond Energy |
| Span | -.94/.88 | -.30/.09 | .27/.08 | .31/.09 |
| Bulge | - | **.07/.00** | -.18/.03 | -.38/.14 |
| Warp | | - | -.58/.34 | **-.03/.00** |
| Skew | | | - | **-.04/.00** |

| Type VIII | | | | |
|---|---|---|---|---|
| | Bulge | Warp | Skew | H-Bond Energy |
| Span | -.64/.40 | **.00/.00** | .12/.01 | .26/.07 |
| Bulge | - | -.66/.44 | -.36/.13 | -.49/.24 |
| Warp | | - | .61/.38 | .49/.24 |
| Skew | | | - | .20/.04 |

| Type IV | | | | |
|---|---|---|---|---|
| | Bulge | Warp | Skew | H-Bond Energy |
| Span | -.58/.33 | .02/.00 | **.01/.00** | .09/.01 |
| Bulge | - | -.63/.40 | .06/.00 | **.02/.00** |
| Warp | | - | .03/.00 | -.07/.01 |
| Skew | | | - | -.14/.02 |

**Table S1. Cross-correlation matrices for turn parameters in the classical types**. Cross-correlation matrices in the classical types for the set of geometric turn parameters and 4>1 H-bond energy. The cell representing each parameter pair in each type contains values for both $r$, the Pearson correlation coefficient for the relationship, and $r^2$, which measures the variance in each member of the pair that can be explained by a best-fit linear model based on the other member, and serves as a metric for the strength of the relationship. Correlations that meet a p-value threshold of .05 for statistical significance (after adjustment for multiple testing using the Bonferroni correction[8]) are set in ordinary type, while those judged not significant are set in bold (most relationships are significant).

The significance of each correlation is evaluated using the procedure described in section 2.2.2 below. The extensive cross-correlation between turn parameters is demonstrated by the fact that of the 90 inter-parameter relationships evaluated between the 4 geometric parameters plus H-bond energy across the 9 classical types, 75 correlations meet the threshold for significance, with most correlations far exceeding that threshold. Most correlations are incapable by themselves of explaining the majority of the variance of the involved parameters (in 53 of the 75 significant relationships, less than a third of the variance in one parameter can be explained by its partner), but they form an interconnected web in which a parameter is influenced, both directly and indirectly, by multiple other parameters.

Key inter-parameter relationships include significant negative correlations between span and bulge in all classical types, with $r$ values ranging from -.58 in type IV to -.94 in type VIb in the type sequence IV→VIII→I'→I→VIa1→II→II'→VIa2→VIb. This result can be rationalized by a simple wire-arch model for a turn: as the span of a fixed-length, semi-rigid wire that forms an arch between two points is reduced, the height of the arch (its bulge) increases.

A fixed-length wire-arch model also predicts a negative correlation between bulge and warp: other factors being equal, as increasing warp brings larger BB excursions in the z dimension above and below the turn plane, the excursion in the x dimension, which corresponds to bulge, should decrease to conserve BB length. Negative correlations do exist between bulge and warp in all types except VIa2 and VIb, with values ranging from -0.35 in type II' to -.78 in type I, in the type sequence II'→II→IV→VIa1→VIII→I'→I.

The relationship between warp and span, however, cannot be explained by conservation of BB length in a wire-arch model, since significant positive correlations exist between span and warp in types I, I', II, II', IV and VIa1, with coefficients ranging up to .55 in the type sequence IV→II'→I'→II→I→VIa1 (although the correlation in type IV is marginal, explaining less than 1% of the variance). This departure from the simple model is not surprising, since bond rotations in actual turns introduce degrees of freedom which complicate the picture by altering the shape of the main-chain path. In type I turns, for example, rotations of the first and third peptide bond planes as warp increases can widen the turn, overcoming pure fixed-length effects.

The strongest correlation involving skew ($r$ = .61), seen in type VIII between skew and warp, occurs because the turn must flatten as it's third residue opens into a more fully extended conformation, and that extension also increases the turn's C-terminal half-span, promoting a more negative skew (see the Ramachandran plots in Figure 3c,d in the main paper).

It should be noted that the correlations shown here are derived for the full ranges of each of the parameters involved, and they may not hold for particular subsets of these ranges. For weaker overall correlations in particular, there may be no correlation in some parts of the parameter ranges, or even correlations opposite to those found in the full ranges. The relationship between two parameters within specific ranges can be evaluated with the ExploreTurns tool, by selecting separate turn sets in higher and lower ranges of one parameter and measuring the difference in the average value of the other parameter between the sets.

#### 2.2.2 Statistical significance of parameter correlations

The significance of each correlation between a pair of parameters is evaluated using the $t$ statistic[9], and a correlation is judged significant if it meets a p-value (pVal) threshold of .05, after correction for multiple testing using the conservative Bonferroni correction[8]. The Bonferroni procedure computes a pVal threshold for the individual correlations which satisfies a desired aggregate pVal' for the set of all correlations, which itself specifies the probability that one or more correlations as high or higher than an observed individual correlation would be seen by chance. The pVal threshold that applies to each individual correlation is computed from the aggregate pVal' as:

$$pVal = 1.0 - (1 - pVal')^{(1/C)}$$

where C is the number of correlations under evaluation. An aggregate pVal' of .05 is chosen, which for C = 90 results in an individual pVal threshold of 5.698E-4, or a one-sided pVal threshold of 2.849E-4.

The $t$ statistic for each correlation is:

$$t = r * \sqrt{(n-2)/(1-r^2)}$$

where $n$ is the number of turns in the classical type. The pVal for each correlation in a classical type is found from $t$ by consulting a table of the t-statistic for $(n-2)$ degrees of freedom (where $n$ is the number of turns in the type), and if this value is less than or equal to the individual pVal threshold then the correlation is judged to be significant.

#### 2.3 Parameters and structure quality

Since geometric parameters enhance the Euclidean-space picture of turn geometry, they may aid in structure validation, by helping to detect and characterize suspicious conformations. Extreme parameter values may in some cases reflect BB stress built up over multiple residues, and parameters could complement existing validation tools not designed to detect these stresses.

Figure S3 investigates the question of whether extreme parameter values are correlated with the occurrence of structural outliers detected by existing methods. The figure presents bar charts of the average per-turn sums of the per-residue counts of geometric quality criteria with outliers, extracted from the PDB's residue-property plots[10]. These counts, which are sampled across the distributions of span, bulge, skew and warp in the global turn set, show clear peaks at the extremes of low span, low bulge, negative and positive skew, and high warp. The span results seem simple to rationalize, since (other factors being equal) turns with the smallest spans have the highest average BB curvature, which is likely associated with more extreme bond angles and more common steric clashes within the turn. A similar argument may explain the increased outlier frequencies at the extremes of skew, warp, and bulge: inspection shows that turns with extreme values of these parameters can exhibit structural contortions, including high curvature, which may result in atypical bond angles and/or bring turn atoms unusually close together.

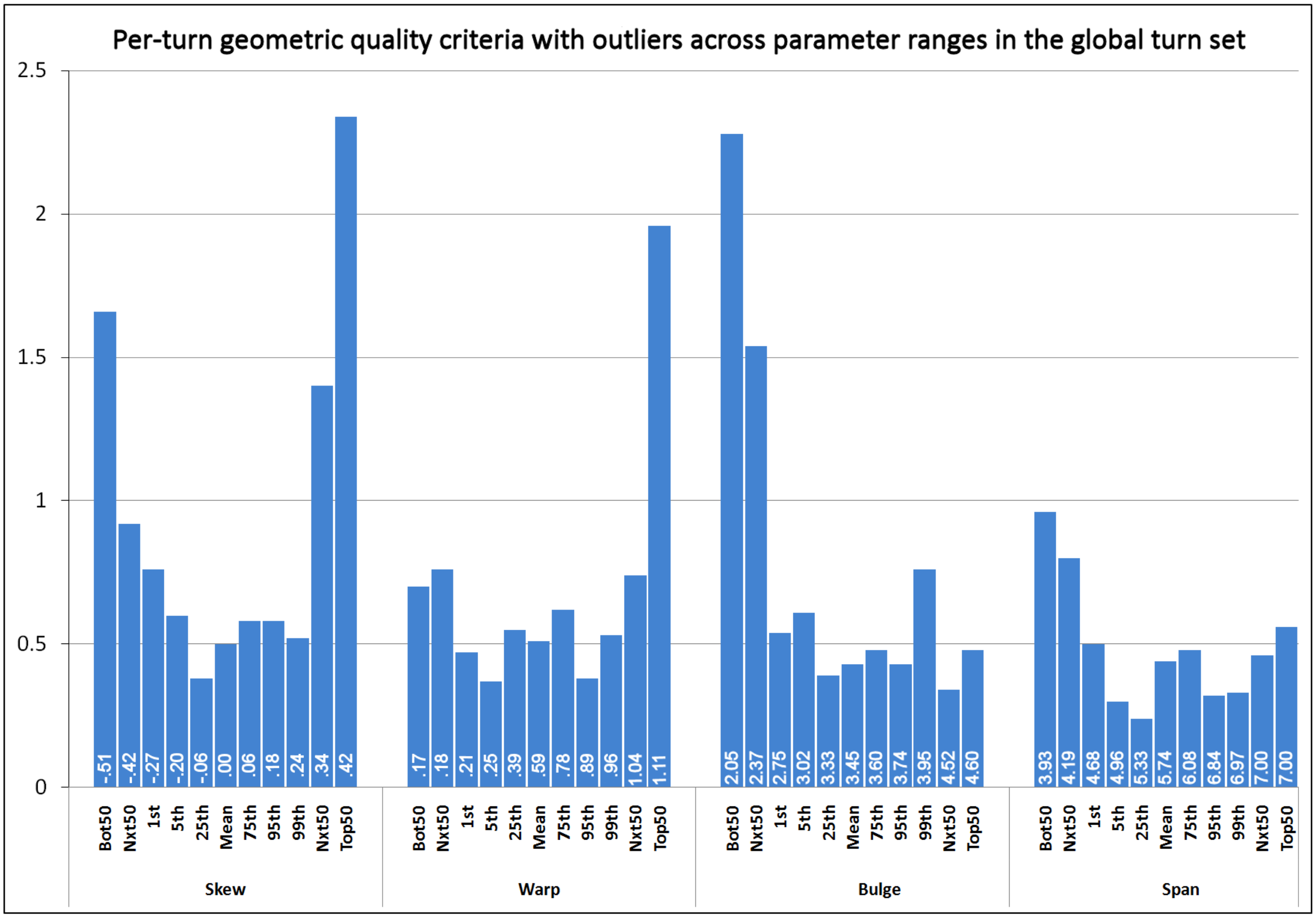


**Figure S3. A PDB outlier metric is commonly elevated in turns with extreme parameter values in the global turn set.** Bar charts of the averages of the per-turn sums of per-residue counts of geometric quality criteria with outliers, plotted across the parameter ranges for skew, warp, bulge and span in the global turn set. Plotted frequencies, computed from data extracted from the PDB's residue-property plots[10], represent lower bounds. Values are computed for 100-turn subsets sampled across the distribution of each parameter, including subsets at the absolute extremes, the 1st, 5th, 25th, 75th, 95th, and 99th percentiles of parameter value, and the mean parameter value. The 100-turn subsets at each extreme are divided into two 50-turn sub-subsets to improve discrimination. Each bar is labelled at its base with the average parameter value in its subset. The plots show clear peaks at the extremes of negative and positive skew, high warp, low bulge and low span, suggesting a relationship between parameter extremes and structure quality that may prove useful in structure validation.

Although Figure S3 shows elevation of the outlier metric at globally extreme parameter values, it does not address the question of whether outliers are higher when parameter values are extreme only for the neighborhood of a turn's particular recurrent BB geometry. Figures S4, S5 and S6, which chart the relationships between parameter extremes and structural outliers for turns partitioned by BB geometry using BB cluster medoids[3], show that the outlier metric is frequently elevated at parameter values that are only locally extreme. The global and geometry-specific results therefore both suggest a relationship between parameter extremity and poor structural quality which may support a role for turn parameters in the identification and characterization of structurally suspicious turn conformations.

The ProfileTurn web tool, in combination with the BB cluster-specific outlier frequency charts presented here, may already provide some utility for structure validation: with ProfileTurn, a user can classify an uploaded turn by BB cluster, measure its geometric parameters, then consult the charts to determine whether any parameter values are structurally suspicious for the turn's cluster.

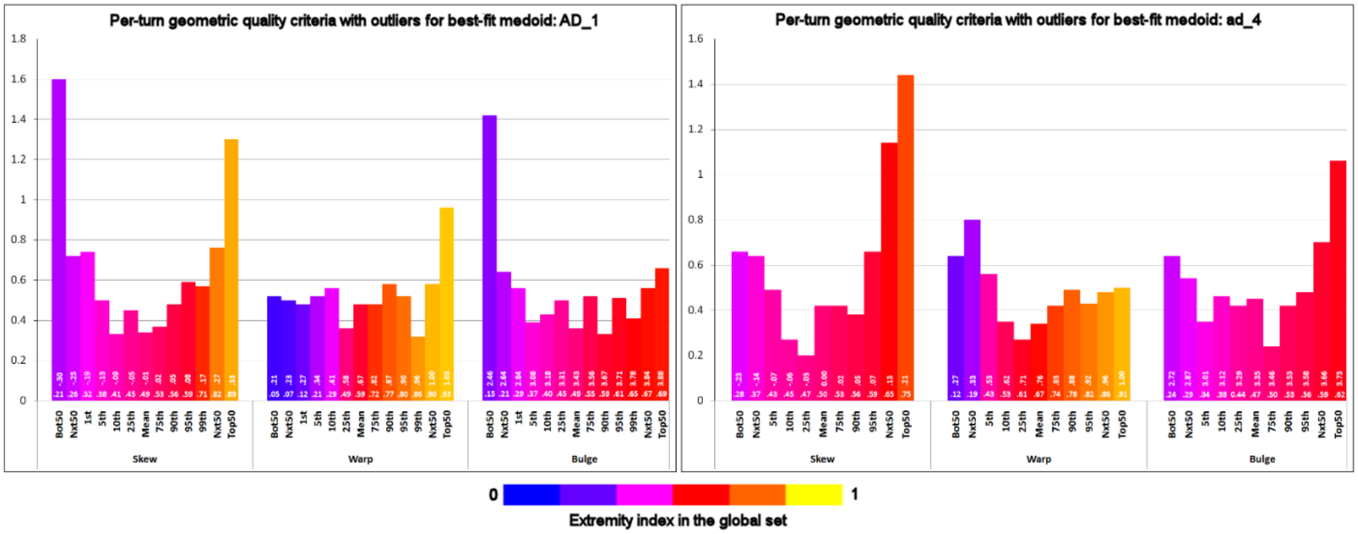


**Figure S4. In the type I/I'-associated turn BB geometries, a PDB outlier metric is commonly elevated at parameter values that are only locally extreme.** Bar charts of the averages of the per-turn sums of per-residue counts of geometric quality criteria with outliers, plotted across parameter ranges for skew, warp and bulge for turns with best-fit BB cluster medoids AD_1 (left) and ad_4 (right), which are associated with classical types I and I' respectively. Plotted values, computed from data extracted from the PDB's residue-property plots[10], represent lower bounds. For type AD_1, frequencies are computed for 100-turn subsets sampled across the distribution of values of each parameter, including subsets at the absolute extremes, the 1st, 5th, 10th, 25th, 75th, 90th, 95th, and 99th percentiles, and the mean value. For the smaller ad_4 cluster, the subsets at the 1st and 99th percentiles are omitted, since these lie very close to the extremes. The 100-turn subsets at each extreme are divided into two 50-turn sub-subsets to improve discrimination. Each bar in a chart is labelled at its base with a global extremity index that indicates where the mean parameter value for the corresponding subset lies in the parameter's distribution in the global turn set, with values ranging from 0, corresponding to the minimum of the global distribution, through .5 for the global mean, to 1 for the global maximum. Bars are color-coded by increasing extremity index according to the legend. Each bar is also labelled, above its extremity index, with the average parameter value in its subset.

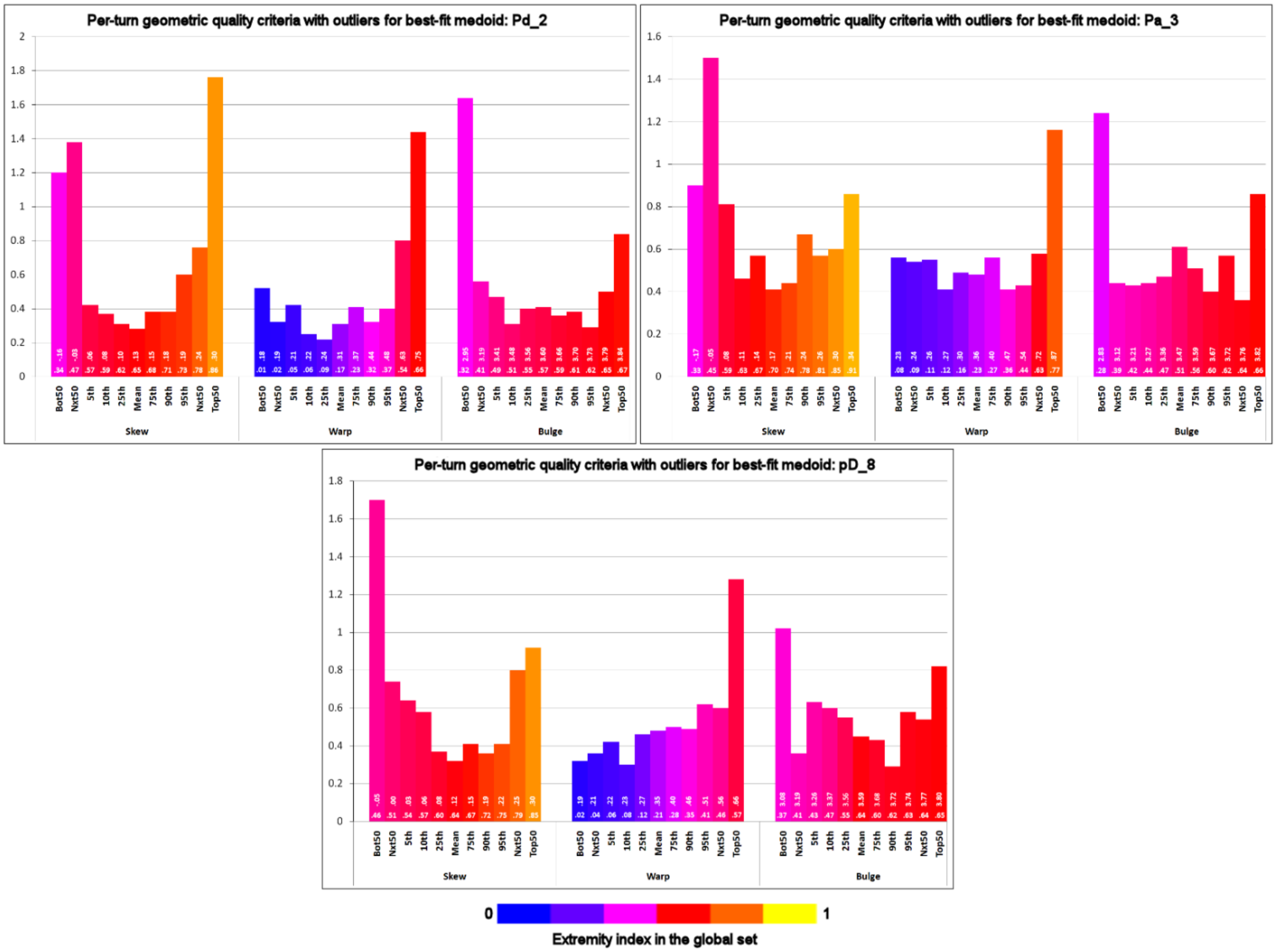


**Figure S5. In the type II/II'-associated turn BB geometries, a PDB outlier metric is commonly elevated at parameter values that are only locally extreme.** Bar charts of the averages of the per-turn sums of per-residue counts of geometric quality criteria with outliers, plotted across parameter ranges for skew, warp and bulge for turns with best-fit BB cluster medoids Pd_2 and Pa_3, associated with classical type II, and pD_8, associated with classical type II'. Chart construction follows the general model of Figure S4.

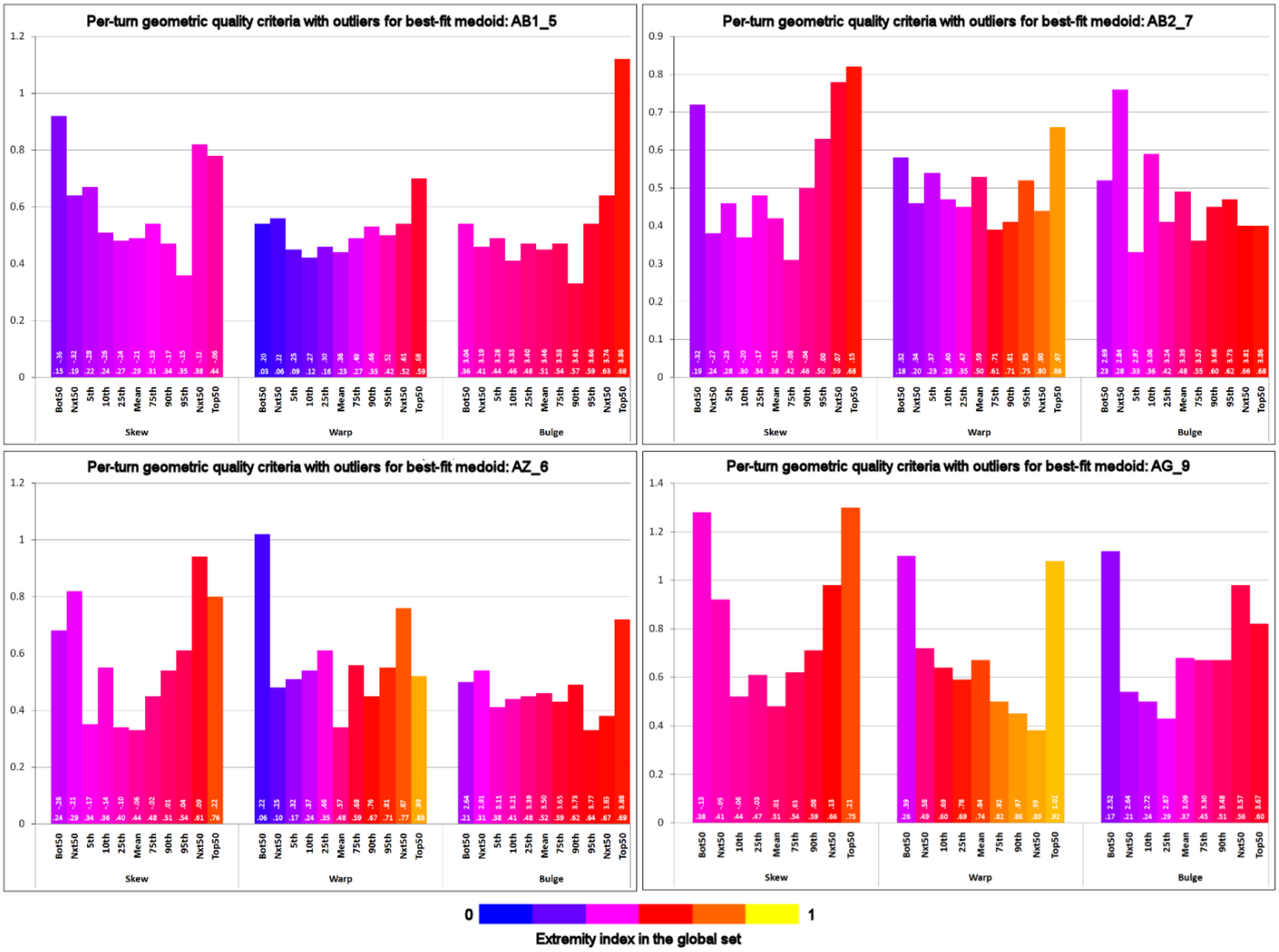


**Figure S6. In the type VIII-associated turn BB geometries, a PDB outlier metric is frequently elevated at parameter values that are only locally extreme.** Bar charts of the averages of the per-turn sums of the per-residue counts of geometric quality criteria with outliers, plotted across parameter ranges for skew, warp and bulge for turns with best-fit BB cluster medoids AB1_5, AZ_6, AB2_7 and AG_9, which are associated with classical type VIII. Chart construction follows the general model of Figure S4.

### 3 | Discussion

#### 3.1 Effect of turn BB geometry on the interpretation and utility of the warp parameter

The differences in BB geometry between the classical turn types or BB clusters affect the interpretation and utility of the warp parameter. Types I and I', with ideal dihedral angles in the helical or near-helical regions at both central residues, are the classical types with the largest average warp, and in these types the BB between $C_{\alpha 1}$ and $C_{\alpha 4}$ commonly executes excursions in the z dimension, above and below the turn plane, which are roughly equivalent in magnitude in each half of the turn, but of opposite sign: in type I, excursions below the plane in the first half of the turn are approximately "balanced" by excursions above the plane in the second half, while in type I' the excursions in each half are reversed.

Type VIII, with ideal angles in the helical region at the first central residue and the strand region at the second, shows a lower average warp than I/I', but the warp distribution covers about the same range as that of type I, and the strand conformation at the second central residue does not in general prevent a rough balance between the BB excursions in the two halves of the turn, when the turn has substantial warp.

Since 76% of higher-warp turns (defined as warp >.6Å) come from types {I, I', VIII}, and 23% come from the structurally unclassified type IV, which contains many turns with conformations close to those of {I, I', VIII}, a large majority of higher-warp turns show an approximate balance between z-dimensional BB excursions in the two turn halves. Furthermore, in types {I, I', VIII}, the identity of the type indicates the signs of the excursions in each half of the turn (-/+ in I and VIII, +/- in I'), so a single warp parameter, taken together with the turn type, provides a useful measure of both the amplitude and sign of the excursions in each turn half (a two-parameter warp measure was tried, but it provided little additional descriptive power).

Warp is less meaningful as a discriminator within types {II, II'}, which comprise 15% of the dataset and exhibit polyproline/strand ideal angles at the first central residue and helical angles at the second, for two reasons. Firstly, these turns show the smallest average warp of all classical types, with many lying close to the limit of flatness, and secondly, the higher-warp turns of each of these types do not exhibit the single, characteristic polarity in their z-dimensional excursions in the turn halves that is seen in types {I, I', VIII}.

In type VI turns {VIa1, VIa2, VIb}, which comprise 1.7% of the dataset, warp is also less meaningful, due to limited warp range in these types. The z-dimensional BB excursions in the types are usually "unbalanced", since the BB proceeds below the turn plane in the first half of the turn and is then commonly directed downwards again in the second half by the central $cis$-peptide bond.
